# Multi-scale spatial transcriptomic atlas of the whole mouse spinal cord

**DOI:** 10.64898/2026.10.24.761397

**Authors:** Guang-Wei Zhang, Jing Li, Junxiang J. Huang, Huizhong Whit Tao, Ke K. Zhang, Li I. Zhang

**Author notes:** Correspondence: Ke K. Zhang; Li I. Zhang. These authors contributed equally to this work (Guang-Wei Zhang, Jing Li, Junxiang J. Huang).

## Abstract

The spinal cord coordinates sensory, motor and autonomic functions through diverse neuronal and non-neuronal populations, yet how molecularly defined cell types are organized across its full extent remains incompletely understood. Here we use Stereo-seq to generate a genome-wide, single-cell-resolution spatial atlas of the whole adult mouse spinal cord. Profiling 322,827 cells across 81 anatomically registered transverse sections spanning cervical, thoracic, lumbar and sacral levels, we resolve 12 major cell classes and 94 transcriptomic subclusters. Major cell classes occupy characteristic anatomical territories conserved along the rostrocaudal axis, whereas individual subclusters exhibit reproducible laminar, mediolateral and segment-associated distributions. Transcriptome-wide analysis identifies coordinated spatial gene-expression modules that delineate anatomical domains independently of predefined cell-type labels. Notably, oligodendrocyte subtypes exhibit a distinct and reproducible laminar organization within spinal cord white matter across spinal levels, revealing finer structural subdivisions beyond conventional anatomical descriptions. Integrating cell-type identities with local spatial organization further defines hierarchical domains that recapitulate canonical anatomy while resolving finer subdivisions within established regions. Together, these findings define a multi-scale cellular architecture linking transcriptomic diversity, spatial gene-expression programs and anatomical organization throughout the spinal cord.

## Introduction

The spinal cord integrates sensory input with motor output and autonomic control through neural circuits arranged in a highly ordered anatomical scaffold [1–3]. Classical studies have defined this scaffold in terms of segmental position, dorsal–ventral lamination, motor columns and surrounding white-matter tracts [4–7], creating the anatomical coordinate system through which spinal circuits are still described and manipulated. These cytoarchitectural landmarks are essential, but they describe tissue shape, cell position and morphology rather than molecular identity. They therefore leave unresolved how transcriptionally defined cell types, their subtype diversification and their local spatial relationships are organized within the intact adult cord.

Single-cell and single-nucleus RNA sequencing have reshaped the molecular taxonomy of the spinal cord and of the brain-wide neurons that project to it [8,9], resolving diverse neuronal, glial and vascular populations and marked heterogeneity within each major class [10–15]. These studies have established robust taxonomies of dorsal-horn sensory interneurons [10,11,16], ventral and cardinal neuron classes [14,17,18], cholinergic and motor populations [13,19–21], and have begun to harmonize cell types across datasets and species [12,15,22,23]. Yet comprehensive profiling of the adult spinal cord remains technically challenging. The tissue is small relative to the brain, regionally heterogeneous and densely myelinated, with long cellular processes that are poorly preserved by dissociation, creating risks of selective cell loss and transcriptional artefacts. Consequently, many atlases rely on nuclei rather than intact cells, and most sample selected levels, regions or populations. Most importantly, dissociation removes native position—the feature on which classical spinal anatomy is built.

Spatial transcriptomics addresses this limitation by measuring gene expression while retaining anatomical context [24,25]. Targeted, probe-based approaches have generated valuable cell-type maps in defined tissues [24,26–28], but they are constrained to genes selected in advance and therefore limit discovery of spatial patterns involving genes outside the selected panel. Genome-wide spatial methods, including DNA-nanoball (DNB) capture arrays, now enable high-resolution profiling across large tissue areas [25,29–33], making it possible to ask how molecular identity is organized across an entire organ [34,35]. However, no genome-wide, single-cell-resolution spatial atlas has yet spanned the whole adult spinal cord, including cervical, thoracic, lumbar and sacral segments. Thus, the spatial logic linking molecular cell types, anatomical domains and genome-wide gene programs across the intact cord remains unknown.

We therefore asked whether the adult mouse spinal cord is organized as a unified, multi-scale molecular architecture in which transcriptional diversification is accompanied by spatial diversification at progressively finer anatomical scales. Using Stereo-seq with single-cell segmentation, we generated an atlas of 322,827 anatomically registered cells spanning cervical, thoracic, lumbar and sacral segments and resolving 12 major cell classes and 94 subtypes. We analysed this organization at three coupled scales: the stereotyped territories of major cell classes, the laminar, mediolateral and segmental niches occupied by molecular subtypes, and co-expression modules formed by spatially variable genes (SVGs) across registered anatomical regions. By integrating these layers with local cellular neighbourhood composition, we define how cell identity, subtype diversity and anatomical domain are jointly arranged across the whole adult spinal cord.

## Results

### A whole-cord spatial atlas of 322,827 anatomically registered cells

To determine how molecular identity is organized in native anatomical space across the entire spinal cord, we performed Stereo-seq on 81 transverse cryosections sampled from individual cervical, thoracic, lumbar and sacral segments of 8-week-old adult mice (Figure 1A; Figure S1).

**Figure 1.**
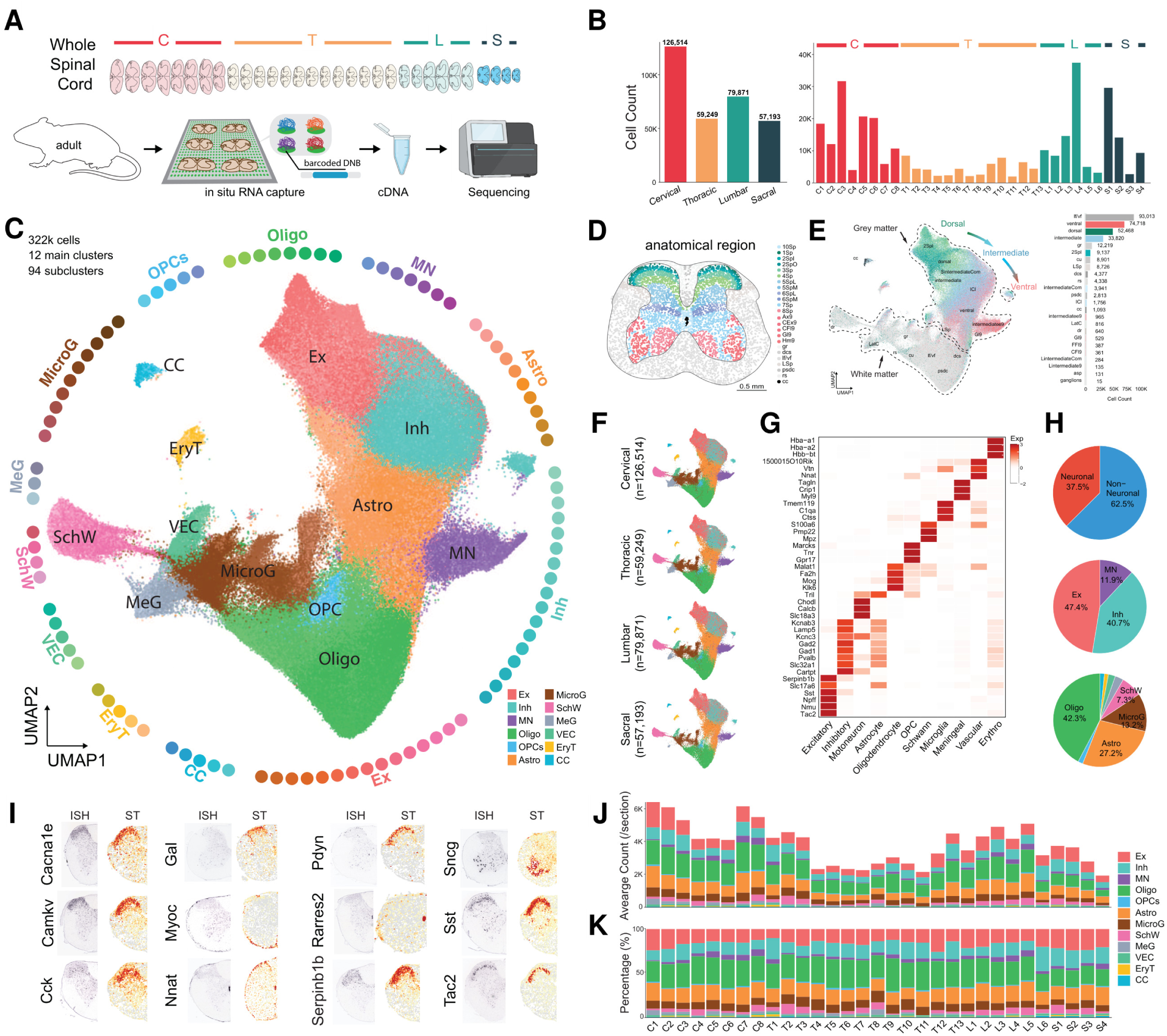
Stereo-seq-based spatial transcriptomic atlas of the adult mouse spinal cord. A. Experimental workflow. Transverse spinal-cord sections spanning individual cervical (C), thoracic (T), lumbar (L) and sacral (S) segments from adult mice were profiled by Stereo-seq, including *in situ* RNA capture on barcoded DNB chips, cDNA library preparation and sequencing. B. Numbers of profiled cells by major spinal level and individual spinal segment. C. UMAP of 322,827 cells resolving 12 major cell classes and 94 subclusters, including excitatory neurons (Ex), inhibitory neurons (Inh), motor neurons (MN) and major non-neuronal populations; peripheral lollipop plots indicate the number of subclusters per class. Astro, astrocytes; Oligo, oligodendrocytes; OPC, oligodendrocyte precursor cells; MicroG, microglia; SchW, Schwann cells; MeG, meningeal cells; VEC, vascular endothelial cells; EryT, erythroid cells; CC, central-canal–associated cells. D. Segment-resolved cross-section with cells assigned to anatomical regions and laminae. Scale bar, 0.5 mm. E. UMAP coloured by registered Allen anatomical region; bar plot (right) shows cells per region. F. Segment-resolved UMAPs showing consistent cell-type organization across major spinal-cord segments. G. Heatmap of marker-gene expression across major cell classes. H. Pie plots of neuronal versus non-neuronal proportions (top) and composition within each compartment (middle, neuronal; bottom, non-neuronal). I. Spatial validation comparing Allen mouse spinal cord in situ hybridization (ISH) and Stereo-seq (ST) expression for representative marker genes. J,K. Mean cell number per section (J) and percentage composition (K) of major cell classes across spinal segments.

Multiple spinal cords were co-embedded in optimal cutting temperature (OCT) compound with defined rostrocaudal offsets, enabling sections from different levels to be captured on the same chip while preserving segmental identity and allowing adjacent Nissl-stained sections to provide anatomical reference (Figure S1). After in situ RNA capture, image-based nuclei detection and Voronoi-based single-cell segmentation, transcripts were assigned to segmented cells and reconstructed as spatially positioned cell bins (Figure S1).

The resulting atlas contained 322,827 high-quality cells with a defined spinal-level assignment, distributed along the rostrocaudal axis (Figure 1F): 126,514 cervical, 59,249 thoracic, 79,871 lumbar and 57,193 sacral cells. Genome-wide expression profiles were highly reproducible across biological and anatomical sources of variation, with strong Pearson correlations for same-segment/cross-animal, cross-segment/same-animal and cross-segment/cross-animal comparisons (Figure S2A–D). Unsupervised integration of all sections with Seurat v5, using SCTransform normalization and Harmony batch correction, resolved 12 major cell classes and 94 subclusters spanning excitatory, inhibitory and motor neurons together with astrocytes, oligodendrocytes, oligodendrocyte precursor cells (OPCs), microglia, Schwann cells, meningeal cells, vascular endothelial cells, erythroid cells and central-canal–associated (ependymal) cells (Figure 1C,G,H). Because each profiled cell retained its spatial coordinates, the dataset could be registered to the Allen Mouse Spinal Cord Atlas [36] and cells assigned to anatomical regions and Rexed laminae [1] (Figure 1D,E). Marker-gene expression was spatially coherent and matched Allen Spinal Cord Atlas *in situ* hybridization (ISH) patterns for representative markers (Figure 1I), and the overall class composition agreed with previous spinal cord single-cell transcriptomic studies [10–14]. In addition, cluster-marker genes (CMGs; the top 2,000 markers for the annotated clusters; Methods) were similarly conserved across the cord: 80.8% were shared among cervical, thoracic, lumbar and sacral sections (Figure S2E), and 93.8% were shared between male and female datasets from L4 and S1 (Figure S2F).

This dataset therefore provides both a molecular taxonomy and an anatomical coordinate framework for analysing cell identity in space. Beyond the expected grey- and white-matter separation, the major organization of the cord was reproducible across segments at transcriptional and spatial levels, and dorsoventral anatomical position corresponded to structure in the integrated UMAP embedding (Figure 1E,F,J,K). We next used this registered whole-cord atlas to test whether spinal cord organization is structured at three coupled scales: major-class territories, subtype-level niches and genome-wide gene programs.

### Major cell classes occupy stereotyped territories conserved across spinal levels

We next asked whether molecularly defined cell classes occupy stereotyped anatomical territories across the whole cord. Representative cervical, thoracic, lumbar and sacral sections showed that each major class had a characteristic spatial distribution: excitatory and inhibitory neurons filled partially overlapping grey-matter domains, motor neurons were concentrated in the ventral horn, oligodendrocyte-lineage cells populated white-matter tracts, and peripheral, vascular and central-canal–associated populations occupied their expected compartments (Figure 2A; Figure S3). Three-dimensional reconstruction of profiled cells illustrated the longitudinal organization of these territories along the rostrocaudal axis (Figure 2A, right). Segment-level maps of excitatory neurons, inhibitory neurons, motor neurons and oligodendrocytes further confirmed that these broad positional relationships were retained from cervical to sacral levels (Figure 2B; Figure S3).

**Figure 2.**
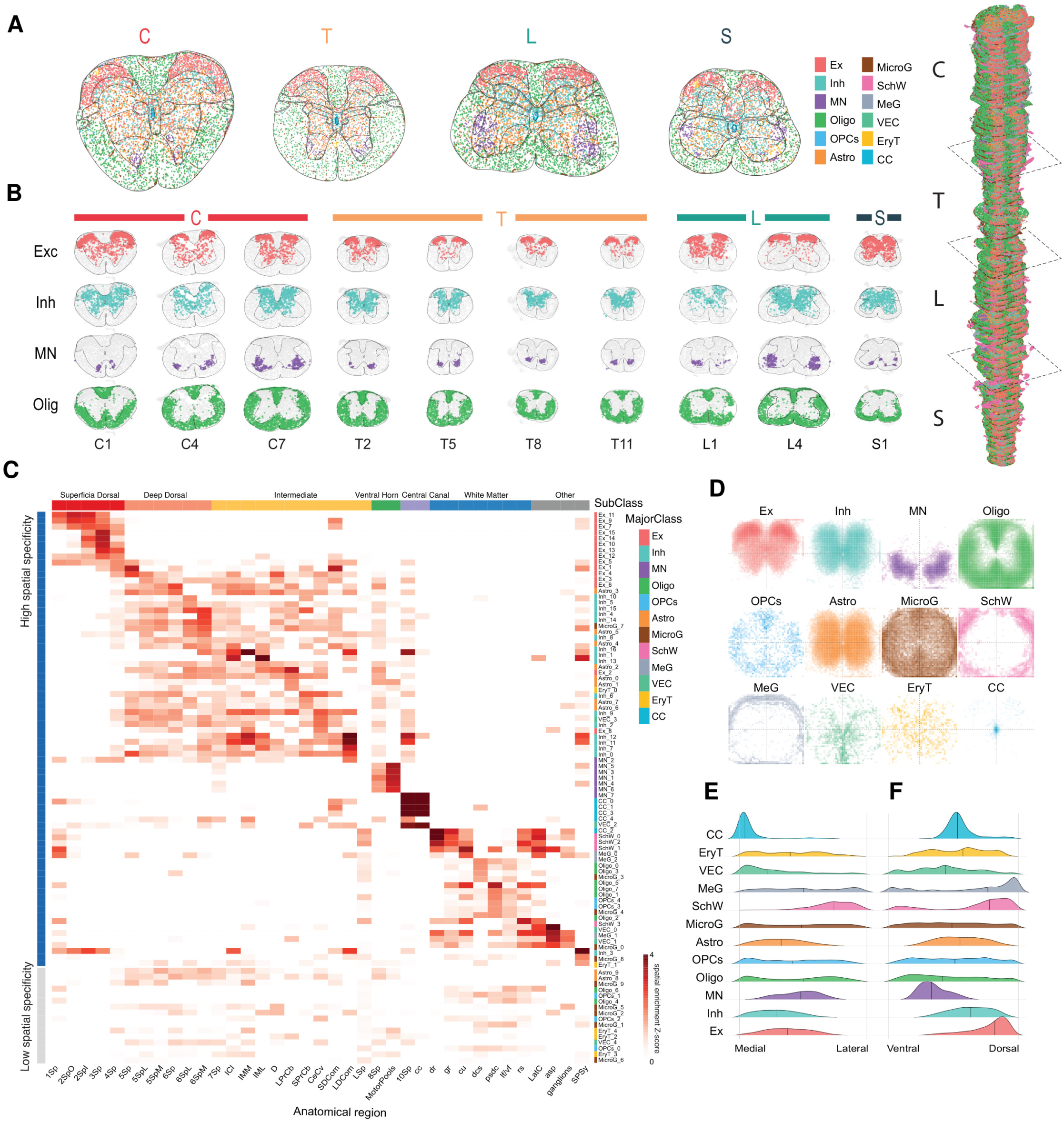
Spatial organization of major cell classes across the adult mouse spinal cord. A. Representative spatial maps of cells in cervical, thoracic, lumbar and sacral sections; each dot is one cell coloured by major class. Right, three-dimensional reconstruction of profiled cells along the rostrocaudal axis. B. Spatial distributions of excitatory neurons, inhibitory neurons, motor neurons and oligodendrocytes across spinal-cord segments (full segment-level distributions in Figure S3). C. Heatmap of subcluster enrichment across anatomical regions; rows are subclusters (left y-axis, spatial-specificity groups defined by spatial autocorrelation; right y-axis, major class), columns are anatomical regions. D. Spatially normalized density distributions of major classes in cross-section. E,F. Ridge plots of each major class along the mediolateral (E) and dorsoventral (F) axes.

We then tested whether this spatial logic also operated below the level of major classes. Spatial-enrichment analysis of all 94 subclusters across registered anatomical regions showed that spatial specificity was not limited to broad lineages, but distinguished finer transcriptomic populations: many subclusters were enriched in one or a few anatomical regions, and subclusters from the same major class frequently occupied distinct territories (Figure 2C). Spatially normalized density maps and mediolateral and dorsoventral ridge plots quantified these class-specific positional preferences and showed that territories are structured along both the mediolateral and dorsoventral axes (Figure 2D–F). Per-segment spatial density profiles further showed that major-class positional distributions were conserved across spinal segments (Figure S3).

Thus, major cell classes define a conserved coarse molecular anatomy, while subcluster enrichment identifies finer spatial niches within classical anatomical domains.

### Genome-wide co-expression modules tile the cord into discrete molecular domains

We next asked whether spatial organization could be detected directly from the genome-wide transcriptome, without using predefined cell-class labels. Using mean log-normalized expression of 21,293 detected genes across 37 registered anatomical regions from all 81 samples, we found that the global expression matrix contained broad anatomical structure, whereas spatially variable genes (SVGs) [37,38] with high spatial specificity revealed sharply region-associated expression patterns (Figure 3A,B). Ranking genes with a spatial-specificity loss-entropy metric (Figure 3B) showed that this organization was carried by a discrete subset of the transcriptome: 5.7% of detected genes (1,212 genes) were high-specificity SVGs and 13.8% (2,941 genes) showed medium specificity, whereas 80.5% (17,138 genes) showed low specificity and were broadly expressed (Figure 3C). Representative gene maps confirmed that SVGs delineated distinct grey-matter, white-matter and laminar domains (Figure 3D).

**Figure 3.**
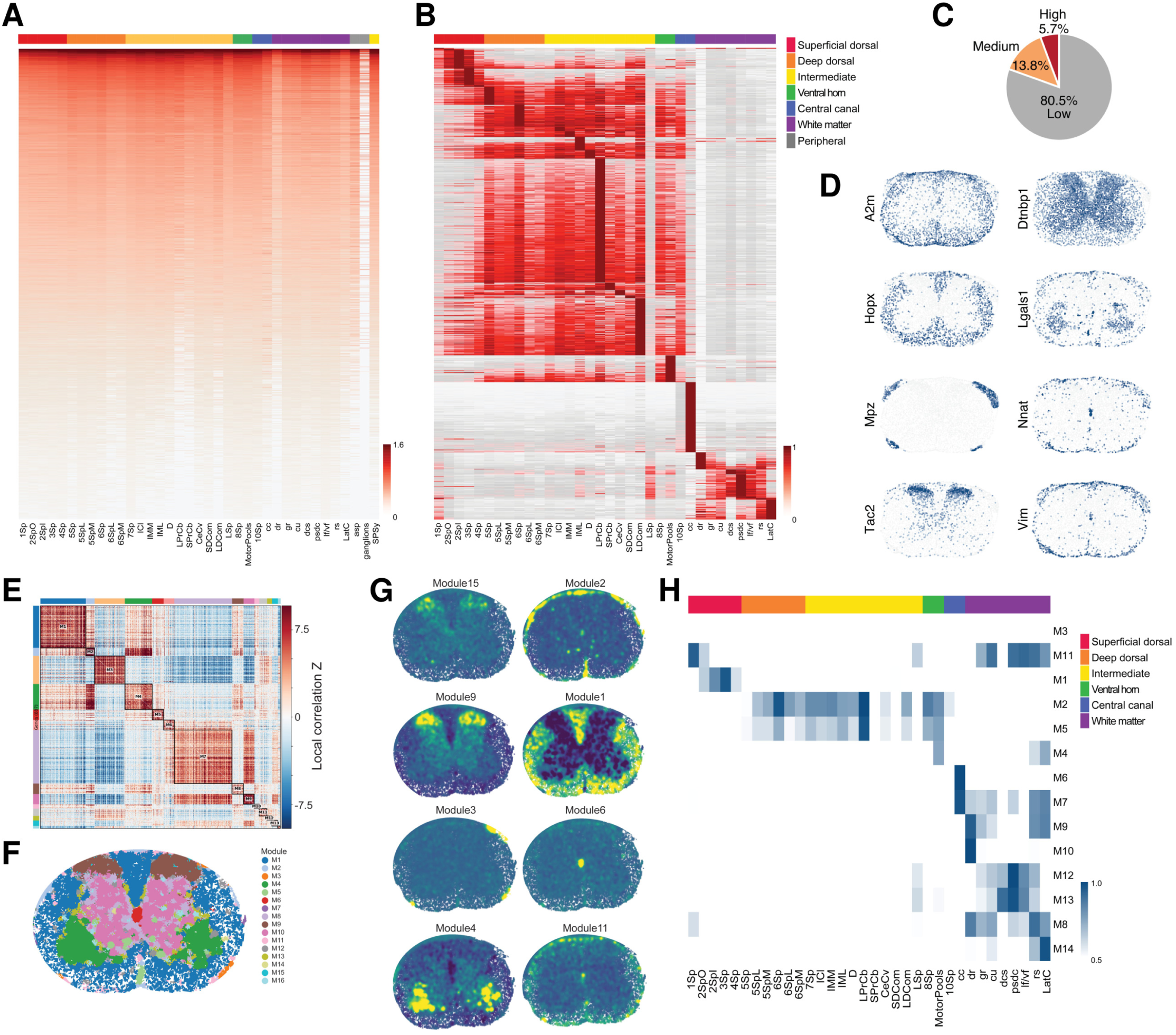
Genome-wide gene-level spatial organization across the adult mouse spinal cord. A. Heatmap of mean log-normalized expression for all detected genes across registered anatomical regions; rows are genes ordered by total expression and columns are regions (81 samples; 21,293 genes across 37 regions). B. Heatmap of spatially variable genes (SVGs) defined by spatial-specificity loss-entropy, ordered to show region-associated expression. C. Pie chart of gene spatial specificity: high (5.7%; 1,212 genes), medium (13.8%; 2,941 genes) and low (80.5%; 17,138 genes). Two of the 21,293 detected genes were not assigned a specificity category. D. Representative spatial expression maps of genes with distinct anatomical distributions. E. Gene–gene local-correlation heatmap defining spatial gene modules; diagonal blocks indicate shared spatial co-expression. F. Spatial module pattern projected onto one representative transverse section. G. Spatial Z-score maps of individual modules in one representative transverse section. H. Heatmap comparing gene spatial modules (rows) with anatomical regions (columns).

We then asked whether these SVGs formed coordinated molecular programs rather than isolated markers. Spatial gene modules inferred from local gene–gene co-expression, using local-correlation analysis, resolved discrete co-expression blocks along the diagonal of the gene–gene correlation matrix (Figure 3E). Projection of module activity onto representative transverse sections showed that individual modules corresponded to coherent anatomical territories (Figure 3F,G). Comparing module activity with registered anatomical regions linked these modules to specific spinal-cord domains—superficial dorsal, deep dorsal, intermediate, ventral horn, central canal and white matter (Figure 3H). Thus, the genome-wide transcriptome encodes a molecular description of anatomical specialization derived without predefined cell-type labels, linking spatial gene expression to tissue-scale anatomy.

### Excitatory neuron subtypes partition classical laminae into conserved dorsoventral niches

We next asked whether transcriptomic diversity within a neuronal lineage is organized into reproducible spatial niches finer than classical laminae. Focused reclustering of excitatory neurons resolved 15 transcriptomically distinct subclusters (Ex_1–Ex_15) with characteristic marker genes (Figure 4A,B). When returned to tissue space, these subclusters grouped into four spatial cluster sets arranged along the dorsoventral axis: G1 (superficial dorsal horn), G2 (upper-mid dorsal horn), G3 (pan-laminar) and G4 (deep dorsal/intermediate) (Figure 4C). These sets were evident across cervical, thoracic, lumbar and sacral segments, and their projection onto a common transverse template and the UMAP embedding showed that spatial position was aligned with transcriptomic structure (Figure 4D–F).

**Figure 4.**
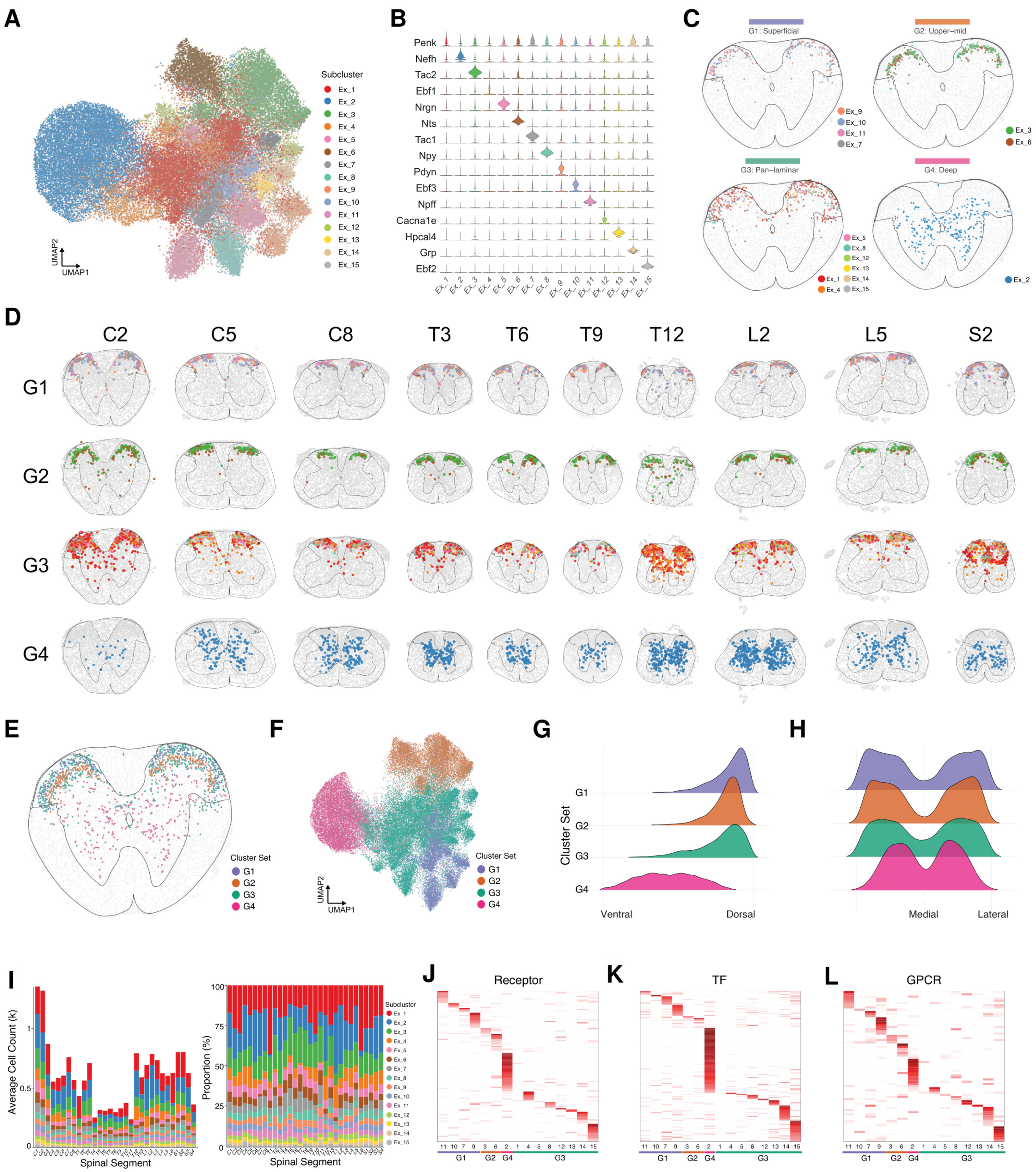
Spatial organization of excitatory neuron subtypes across the whole mouse spinal cord. A. UMAP of excitatory neurons coloured by subcluster (Ex_1–Ex_15). B. Violin plots of representative markers across subclusters. C. Representative transverse sections showing four spatial cluster sets: G1 superficial, G2 upper-mid, G3 pan-laminar, G4 deep. D. Spatial distributions of the cluster sets across representative spinal segments. E. Representative transverse section showing the combined distribution of cluster sets. F. UMAP coloured by spatial cluster set. G,H. Ridge plots of cluster sets along the normalized dorsoventral (G) and mediolateral (H) axes. I. Average cell count (left) and proportional composition (right) of subclusters across segments, showing rostrocaudal enrichment with the largest numbers at cervical and lumbar enlargements. J–L. Heatmaps of receptor (J), transcription-factor (K) and GPCR (L) gene expression across subclusters grouped by spatial cluster set. Source data for the heatmaps in J–L, and for the corresponding heatmaps in Figures S4G and S5G–I, are provided in Supplementary Table 1.

These excitatory neuron niches were conserved across spinal levels but varied locally in abundance and extent, consistent with a stable laminar scaffold overlaid by segment-specific organization. Dorsoventral and mediolateral ridge plots quantified the positional separation of the four cluster sets (Figure 4G,H), and segment-resolved composition analysis showed how subcluster abundance shifted along the rostrocaudal axis (Figure 4I). Finally, group-enriched expression of neurotransmitter-receptor, transcription-factor (TF) and G-protein–coupled receptor (GPCR) genes showed that spatially segregated excitatory populations carry distinct signalling and regulatory profiles, linking anatomical position to molecular specialization (Figure 4J–L).

### Inhibitory neuron subtypes show spatially restricted and segment-spanning organization

We next tested whether inhibitory neurons follow the same subtype–niche principle. Reclustering of inhibitory neurons resolved 17 subclusters (In_1–In_17) with distinct molecular signatures spanning neuropeptides, membrane proteins, calcium-channel genes, receptors, transcription factors and GPCRs (Figure S4A,B,G). Although inhibitory neurons as a class extended through dorsal, intermediate and ventral grey matter, individual subclusters occupied restricted territories that were preserved across levels (Figure S4C,D). Mapping selected groups —In_1, In_2, In_3, In_4 and In_17—across cervical, thoracic, lumbar and sacral segments confirmed that these subtypes formed reproducible domains along the whole cord (Figure S4D; Figure S6).

Positional density analysis provided a clear example of subtype-specific spatial specialization: In_17 formed a distinct superficial-dorsal domain separable along the dorsoventral axis from the remaining inhibitory subclusters (Figure S4E). Segment-level summaries showed that inhibitory subtype composition varied along the rostrocaudal axis while the overall spatial organization of inhibitory populations was retained (Figure S4F). Receptor, transcription-factor and GPCR heatmaps further defined molecular signatures that link inhibitory neuron diversity to functional gene repertoires (Figure S4G).

### Motor neuron subtypes form molecularly and segmentally distinct domains in the ventral horn

Motor neurons provided a complementary test of subtype-level spatial organization because their arrangement is defined by both ventral-horn position and segmental motor architecture. Reclustering of Slc5a7-, Prph- and Chodl-expressing cells resolved motor-neuron subclusters (MN_0–MN_6), each marked by characteristic combinations of canonical motor-neuron identity genes and subtype-specific markers (Figure S5A,B). When projected into tissue space, all motor-neuron subtypes remained ventral but separated by both rostrocaudal and transverse position, forming distinct domains within the ventral horn (Figure S5C,D).

Segment-resolved mapping revealed clear rostrocaudal structure. MN_0 was enriched at cervical and lumbar enlargement segments, where limb-related motor circuitry is concentrated and the largest absolute motor-neuron numbers are found, whereas other subtypes showed complementary or less distinct level-biased distributions (Figure S5D,E; Figure S6). Dorsoventral and mediolateral ridge plots captured the positional organization of subclusters within the ventral horn (Figure S5F), and transmitter-receptor, transcription-factor and GPCR heatmaps identified molecular signatures that distinguished the motor-neuron subtypes (Figure S5G–I). Across excitatory, inhibitory and motor lineages, transcriptional subtypes therefore mapped onto reproducible spatial niches within classical anatomical domains, with segment-specific modulation overlaid on a conserved scaffold.

### Non-neuronal lineages compartmentalize white matter into molecular subdomains

We next asked whether the subtype–niche principle extends to non-neuronal lineages and whether it reveals molecular structure within white matter, whose heterogeneity is often obscured by tract-level anatomy. The non-neuronal compartment resolved astrocytes, oligodendrocytes, OPCs, microglia, Schwann cells, meningeal cells, vascular endothelial cells and erythroid cells, with broadly conserved class composition across levels (Figure 5A–C; Figures S7 and S8). Each class occupied a characteristic compartment: astrocytes [39,40] were concentrated in grey matter, oligodendrocyte-lineage cells concentrated in white-matter tracts [41,42], microglia tiled white and grey matter [43–45], Schwann and meningeal cells occupied peripheral territories, vascular-associated cells followed the vasculature [46], and central-canal– associated cells remained pericentral (Figure 5D–F).

**Figure 5.**
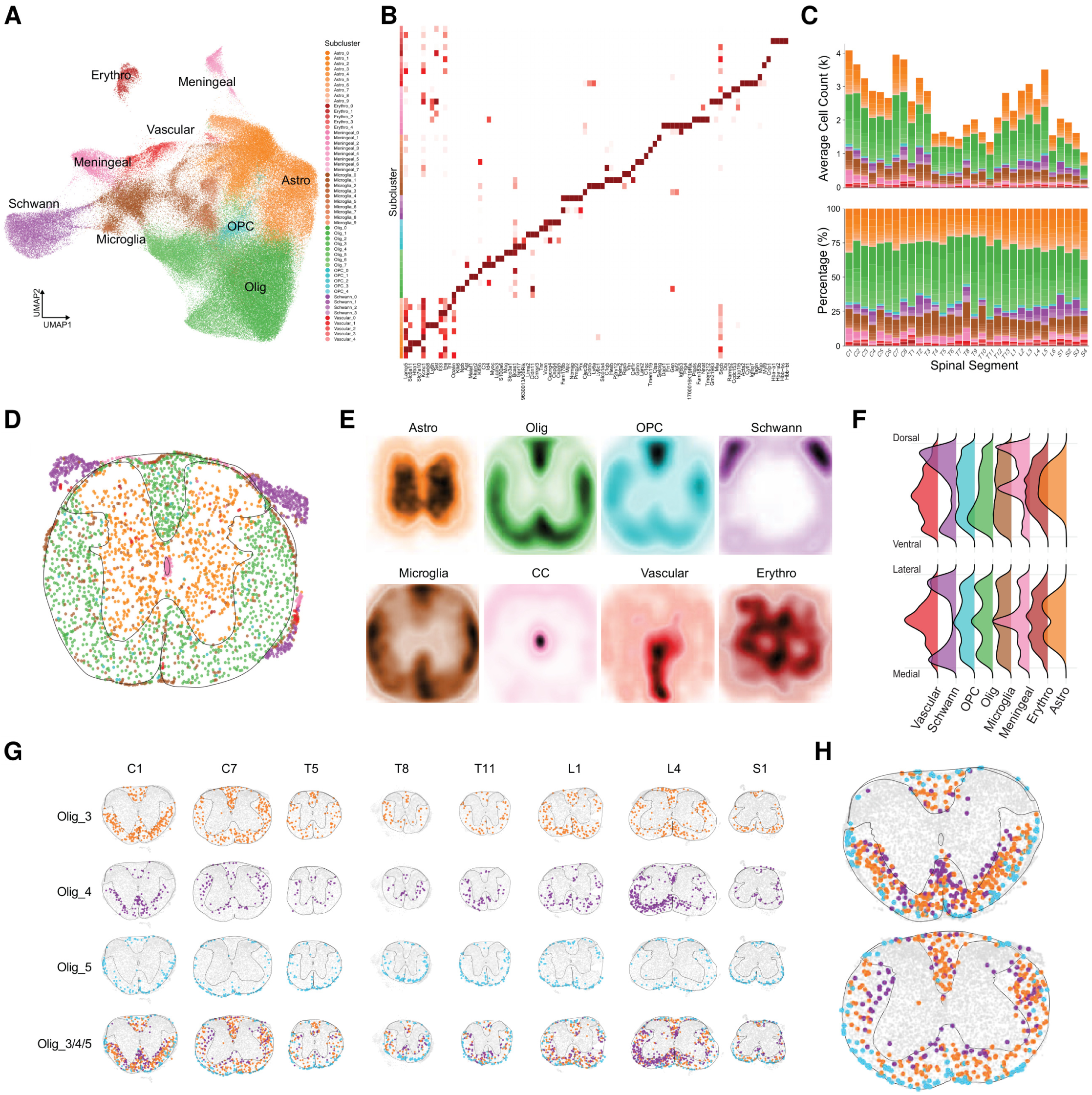
Spatial organization of non-neuronal cell types across the whole mouse spinal cord. A. UMAP of non-neuronal cells coloured by subclass (astrocytes, oligodendrocytes, OPCs, Schwann cells, microglia, meningeal cells, vascular cells, erythroid cells). B. Heatmap of marker-gene expression (column) across non-neuronal subclasses (row). C. Average cell count (upper) and proportional composition (lower) of non-neuronal subclasses across segments. D. Representative transverse section coloured by non-neuronal subclass. E. Spatially normalized density distributions of major non-neuronal types in cross-section. F. Ridge plots of dorsoventral and mediolateral distributions of major non-neuronal types. G,H. Spatial distributions of oligodendrocyte subgroups showing laminar organization within white matter across segments (G) and in representative transverse sections (H).

The clearest subtype-level structure emerged within the oligodendrocyte lineage. Distinct oligodendrocyte subclusters (Olig_3, Olig_4, Olig_5) preferentially occupied different white-matter territories and segregated reproducibly across cervical, thoracic, lumbar and sacral levels, revealing a previously unidentified, reproducible laminar organization of oligodendrocyte subtypes within white matter (Figure 5G,H; Figures S7 and S8). Whereas regional, tract-biased oligodendrocyte heterogeneity has been reported previously [47,48], our genome-wide, whole-cord single-cell data resolve a finer, depth-organized laminar arrangement across spinal levels without a preselected marker panel. Thus, non-neuronal lineages show a similar multi-scale organization to neurons: major classes define broad compartments, and subtypes partition those compartments into finer molecular territories—including within white matter, a compartment often treated as anatomically uniform.

### A hierarchical cell-type–context spatial atlas recapitulates and refines canonical anatomy

Finally, we integrated molecular identity with local spatial context by building a spatial atlas from cellular neighbourhood composition rather than from gene expression directly. For each cell, neighbourhood composition was encoded as a vector, and these vectors were used for dimensionality reduction and clustering. Coarse Level 1 modules captured broad spatial domains —dorsal, intermediate, ventral, central-canal, white-matter and peripheral—as shown by UMAP, representative transverse sections and three-dimensional reconstruction from C1 to S4, which illustrated the longitudinal distribution of these domains (Figure 6A–C).

**Figure 6.**
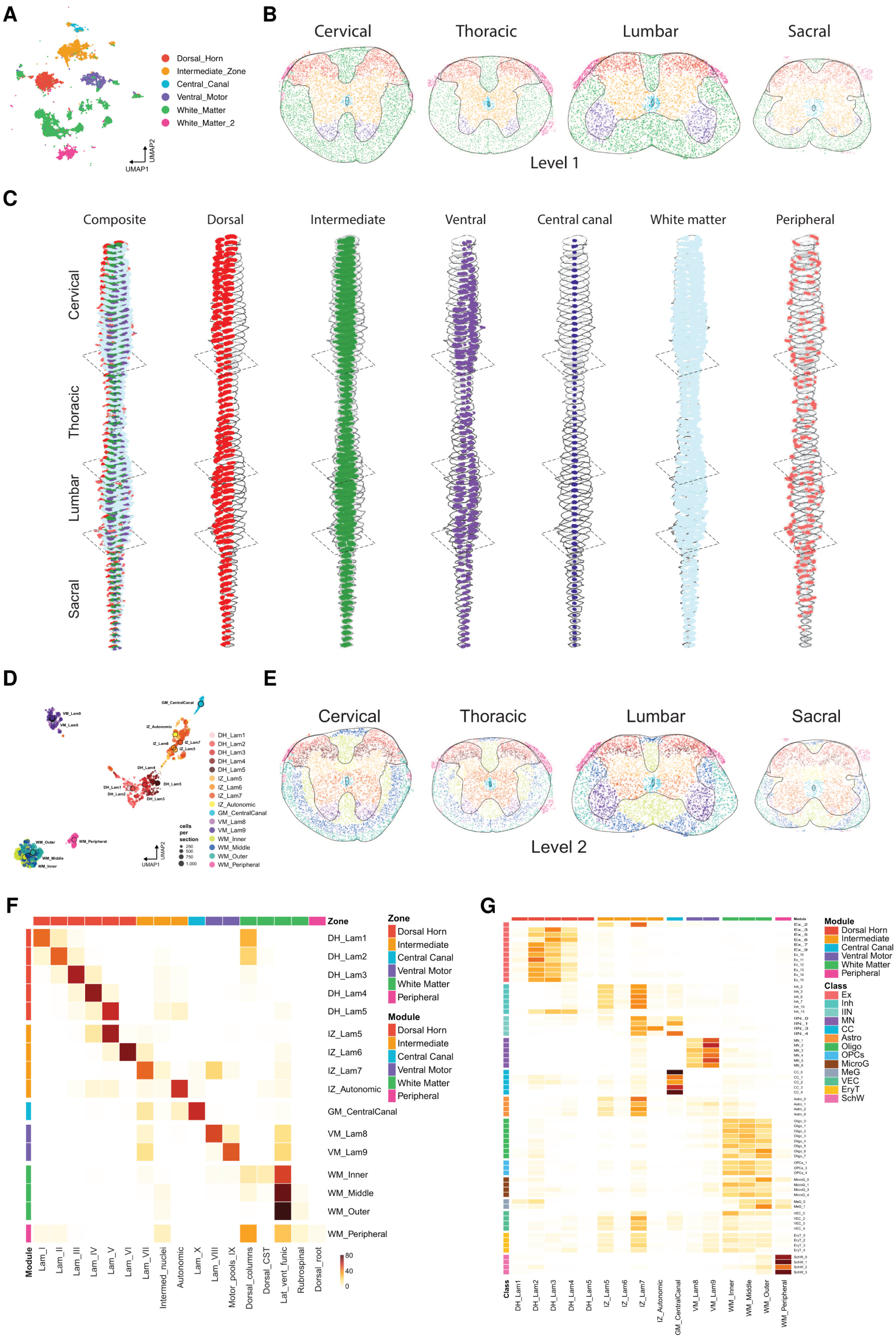
Cell-type–context spatial atlas of the whole mouse spinal cord. A. UMAP of cells clustered by neighbourhood cell-type composition vectors, defining Level 1 spatial-atlas modules. B. Representative C/T/L/S transverse sections coloured by Level 1 module. C. Three-dimensional reconstruction of Level 1 modules along the rostrocaudal axis from C1 to S4, for all modules (left) and for individual modules. D. UMAP of Level 2 modules showing finer anatomical subdivision. E. Representative transverse sections coloured by Level 2 module. F. Heatmap comparing Level 2 modules with canonical anatomical zones; values are the row percentage of each module assigned to each zone. G. Heatmap of the distribution of cell subclusters across Level 2 modules; values are the percentage of each subcluster assigned to each module.

Refined Level 2 modules subdivided these broad domains into finer anatomical patterns (Figure 6D,E). Comparison with canonical anatomical zones showed that Level 2 modules aligned closely with dorsal-horn laminae, the intermediate zone, the central canal, ventral motor columns, white-matter tracts and peripheral territories, while also resolving substructure within them (Figure 6F). Mapping transcriptomic subclusters onto Level 2 modules showed that cell types and spatial domains are tightly linked but not one-to-one: most subclusters contributed to one or a few modules, and each module contained a characteristic combination of cell types (Figure 6G). Together, these analyses establish a hierarchical whole-cord framework in which gene expression, molecular cell identity, anatomical registration and local neighbourhood composition are integrated at single-cell resolution: major classes set coarse territories, subtypes partition them into reproducible niches, genome-wide modules supply the molecular substrate, and integrated spatial modules recapitulate classical cytoarchitecture while resolving molecular subdomains, including within white matter, that cytoarchitecture alone cannot see.

## Discussion

Our results show that the adult mouse spinal cord is organized through coupled molecular and spatial layers. Across 322,827 anatomically registered cells spanning every cervical, thoracic, lumbar and sacral segment, major neuronal and non-neuronal classes occupied stereotyped territories, molecular subtypes subdivided these territories into reproducible niches, and genome-wide co-expression modules partitioned the cord into discrete domains without using predefined cell-type labels. Clustering cells by local cell-type composition provided a complementary spatial-atlas framework that recapitulates canonical anatomy while revealing molecular subdomains not apparent from cytoarchitecture alone. Thus, spinal cord organization is not simply a laminar scaffold populated by cell types; it is a coordinated architecture in which cell identity, subtype diversification and tissue domain are jointly arranged.

### A hierarchical molecular architecture refines classical anatomy

Classical spinal cord anatomy is defined by cytoarchitecture, laminar organization and tract morphology. Our data show that these landmarks correspond to molecularly distinct compartments: major classes define conserved territories, whereas data-driven spatial modules subdivide dorsal, intermediate, ventral, central-canal, peripheral and white-matter domains at finer resolution. This molecular view does not replace classical anatomy, but adds a genome-wide coordinate layer to it. The relationship was most evident when transcriptomic subtypes were returned to tissue space: excitatory, inhibitory and motor subtypes formed reproducible laminar, mediolateral, ventral-horn and segment-biased niches, linking subtype identity to spinal anatomy.

### Molecular compartments within white matter

Within white matter, oligodendrocyte subtypes (Olig_3, Olig_4, Olig_5) occupied a previously unidentified, reproducible laminar arrangement, segregating consistently along the rostrocaudal axis. Regionally biased oligodendrocyte populations have been described in the mouse spinal cord using targeted spatial methods, where mature oligodendrocyte subtypes show tract-level preferences [47,48]. Our genome-wide, whole-cord, single-cell data resolve a finer, depth-organized laminar arrangement that tract-level descriptions did not capture: the compartments are resolved without a preselected marker panel, defined by the full transcriptome of each subtype, and shown to be reproducible across every cervical, thoracic, lumbar and sacral segment. An important open question is whether Olig_3– Olig_5 correspond to previously defined mature oligodendrocyte subtypes, and whether these compartments reflect tract identity, axonal properties or developmental origin—hypotheses now directly testable within this framework. Firm cell-intrinsic conclusions will require *in situ* validation of the distinguishing markers in an independent cohort.

### Subtype niches extend transcriptomic cell-type atlases

Single-cell, single-nucleus and spatial studies have substantially advanced the molecular classification of spinal cord cell types, particularly within dorsal-horn circuits and at lumbar levels [10–15,49]. Our taxonomy is broadly consistent with these classifications, supporting the robustness of major spinal cord classes and subtypes across studies and platforms. Its main extension is anatomical: molecular populations can now be placed within a whole-cord coordinate frame, assessed for laminar and segmental reproducibility, and compared with genome-wide spatial programs. This changes the interpretation of transcriptomic clusters. Excitatory, inhibitory, motor and non-neuronal subtypes are not only molecular groups inferred from dissociated data, but populations assigned to defined tissue domains, allowing cell types, subtype niches and anatomical regions to be interpreted together. Relative to dissociation-based atlases, which discard native coordinates, and to targeted spatial methods, which read a predefined gene panel, untargeted whole-cord profiling adds two capabilities in particular: mapping subtype distributions across spinal levels and identifying coordinated gene expression beyond a predefined marker panel.

### Genome-wide spatial programs as a molecular substrate

A defining strength of genome-wide spatial transcriptomics is that it reveals organization beyond predefined markers or selected cell populations. Our co-expression-module analysis showed that regional structure is encoded not only by annotated cell-type distributions, but also by coordinated gene programs that tile the tissue into dorsal, intermediate, ventral, central-canal, peripheral and white-matter domains. These modules provide the molecular substrate for the spatial framework, linking cell-type architecture to tissue-scale anatomy and explaining why a shared organizational logic is visible from individual subtypes to whole-cord domains.

### Limitations and outlook

The present dataset is transcriptomic and positional; it does not directly report synaptic connectivity, electrical activity or behavioural function, and should therefore be interpreted as a molecular and spatial framework rather than a functional circuit model. In particular, Voronoi-based segmentation assigns transcripts to spatially contiguous bins, so a cell’s measured profile can be influenced by neighbouring cells and ambient RNA; spatially organized results, including the white-matter oligodendrocyte compartments, therefore require orthogonal in situ validation before firm cell-intrinsic conclusions are drawn. Because regional expression partly reflects local cell-type composition, the genome-wide co-expression modules should likewise be read as a description of spatial gene expression rather than as evidence of cell-type-independent regulation. The dataset is also restricted to mouse, so whether the same compartments and niches are conserved in human or non-human primate spinal cord [22,33,50] remains untested. Cell segmentation and anatomical registration introduce further technical constraints that will be refined as spatial methods mature. The value of whole-cord, genome-wide spatial profiling is that it places molecularly defined cell populations and coordinated gene expression along the full rostrocaudal axis. Integrating this framework with circuit-level recordings, perturbation, developmental time courses, and injury and disease models is a natural next step [51–55]. By unifying cell identity, subtype-level niches, genome-wide co-expression modules and anatomically registered spatial modules, this atlas links classical spinal cord anatomy to molecular organization and establishes a framework for studying spinal function, development, injury and disease with molecularly defined cells and gene expression anchored in anatomical space.

## Methods

### Ethical compliance

All procedures followed the Guide for the Care and Use of Laboratory Animals (NIH) and were approved by the Institutional Animal Care and Use Committee (IACUC) of the University of Southern California.

### Animals and housing

Adult (8-week-old) C57BL/6J mice (The Jackson Laboratory; RRID:IMSR_JAX:000664) of both sexes were used (30 males and 9 females). Mice were group-housed with same-sex conspecifics under a 12-h light/dark cycle (lights off at 18:00) with ad libitum access to food and water until non-survival spinal cord extraction.

### Sample preparation

#### Spinal cord dissection and sample embedding

Mice were deeply anaesthetized with isoflurane (5%) and transcardially perfused with ice-cold RNase-free 1× PBS containing 0.5% sodium nitrite. Whole spinal cords were dissected, coated with ice-cold O.C.T. (Tissue-Tek, Sakura Finetek), transferred to embedding molds (Peel-A-Way, 18646D-1) and frozen on dry ice. To maximize use of the Stereo-seq capture surface while preserving segmental information, multiple spinal cords were co-embedded in a single mold with defined rostrocaudal offsets, allowing transverse sections from different spinal levels to be captured together. Samples were stored at −80 °C until sectioning.

The sampled spinal segments and corresponding mouse IDs were as follows: C1, from mouse#3, 9A, 12C; C2, from mouse#9A, 9B; C3, 9B, 9C, 15A twice, 15B, 15C; C4, from mouse#12D; C5, from mouse#9C, 12A, 15A, 15C, 15D; C6, from mouse#9D twice, 15B twice, 15C; C7, from mouse#9B; C8, 9C, 12B; T1, from mouse#15A, 15D; T2, from mouse#9A; T3, from mouse#9D; T4, from mouse#13D; T5, from mouse#13C; T6, from mouse#3, 13A; T7, from mouse#13B; T8, from mouse#13D; T9, from mouse#13C, 15B; T10, from mouse#13A, 15A, 15C; T11, from mouse#13B; T12, from mouse#11A twice; T13, from mouse#16B; L1, from mouse#3, 11C, 13D; L2, from mouse#11C, 13C; L3, from mouse#13A, 16A, 16E; L4, from mouse#11B, 13B, 16C, 16D, 16F, 17A, 17B, 17C, 17D; L5, from mouse#11B; L6, from mouse#16I; S1, from mouse#11D, 13D, 16G, 17E, 17F, 17G, 17H, 17I; S2, from mouse#11D, 13C, 15C, 16H; S3, from mouse#13A; S4, from mouse#3, 13B, 15A, 15B, 15D. Mouse IDs comprise a tissue-block ID and a letter identifying each mouse within the block. Mice #17A–17I were female; the remaining mice were male.

### Data acquisition

#### Stereo-seq library preparation and sequencing

Stereo-seq (SpaTial Enhanced REsolution Omics-sequencing) was performed essentially as described [25,29]. Samples were thawed at −20 °C for ≥30 min and cryosectioned transversely at 10 µm on a Leica CM1950 cryostat. Stereo-seq capture chips (∼20 mm^2^ capture area) were washed twice with nuclease-free water and dried at 37 °C. Sections were mounted onto pre-chilled chips, incubated at 37 °C for 3 min, and fixed in pre-cooled methanol at −20 °C for 30 min. After fixation, sections were stained with Qubit ssDNA reagent (Thermo Fisher, Q10212) in 5× SSC for 5 min, washed in 0.1× wash buffer (0.1× SSC with 5% RNase inhibitor, BGI-Research 1000028496), dried and imaged on a Motic PA53 FS6 microscope (FITC channel, 10× objective) to record nuclei and tissue morphology. Tissue was permeabilized with PR enzyme (BGI-Research) in 0.01 M HCl at 37 °C for 12–15 min. Captured mRNA was reverse-transcribed in situ at 42 °C for 3 h. Tissue was removed (55 °C, 10 min), and cDNA was released at 55 **°**C overnight, purified with VAHTS DNA clean beads (0.8×, Vazyme N411-03) and amplified (cDNA primer 5′-CTGCTGACGTACTGAGAGGC-3′; 95 °C 5 min; 15 cycles of 98 °C 20 s, 58 °C 20 s, 72 °C 3 min; 72 °C 5 min). Amplified cDNA was purified (0.6× beads), quantified (Qubit dsDNA Assay, Thermo Q32854), and 20 ng was fragmented with in-house Tn5 at 55 °C for 10 min and stopped with 0.02% SDS. Fragmented products were amplified, purified with AMPure XP beads (0.63× and 0.15×), used for DNA-nanoball (DNB) generation and sequenced on an MGI DNBSEQ-Tx.

#### Raw data processing

DNBSEQ-Tx reads were processed with the STOmics Analysis Workflow (SAW). Read 1 contained the coordinate identity (CID, 1–25 bp) and molecular identity (MID, 26–35 bp), and Read 2 contained the cDNA sequence. CIDs were matched to chip barcodes to recover spatial position, valid reads were filtered, and clean reads were aligned to the GRCm38 mouse reference genome [56]. Reads overlapping exons by >50% were counted as exonic; reads overlapping adjacent introns were annotated as intronic, and all remaining reads as intergenic. Exonic reads were used to build a CID-indexed expression matrix, with duplicate MIDs removed and MID-corrected gene counts computed. The Stereo-seq DNB array has a nanoball diameter of ∼220 nm and a centre-to-centre spacing of 500 nm.

#### Nissl staining and segment confirmation

Sections adjacent to those used for Stereo-seq were stained with NeuroTrace (435/455 Blue Fluorescent Nissl Stain, N21479) and imaged on a Zeiss system. Briefly, fresh-frozen sections were fixed in 4% paraformaldehyde in 1× PBS (5 min, RT), rinsed in PBS, stained with 1% NeuroTrace in 0.1% Triton X-100/PBS (5 min, RT), washed and coverslipped.

### Data processing

#### Cell segmentation

Segmentation was performed in three steps. First, we identified the coordinates of cellular nuclei from the ssDNA nuclear-staining images. To achieve robust nucleus detection and segmentation, distance-transform watershed instance segmentation and size filtering were combined with ilastik-based pixel classification [57]. This strategy enabled effective background removal while ensuring that the identified objects were consistent with expected nuclear morphology.

Second, we transformed the coordinates of nuclei identified from the ssDNA images into the coordinate system of the sequencing data by aligning the ssDNA images with the UMI spatial maps using a set of reference keypoints. UMI spatial maps were reconstructed directly from bin1 sequencing data as greyscale images, with pixel intensity corresponding to the total UMI count at each DNA nanoball (DNB) location. The Stereo-seq capture chips contain a spaced grid pattern that is visible in both the ssDNA images and UMI spatial maps after appropriate adjustment of brightness and contrast. Grid intersections were identified as reference keypoints and used to estimate the geometric transformation between the two imaging modalities. The resulting transformation was then applied to map the ssDNA-image-derived nuclear coordinates onto the sequencing-chip coordinate system.

Finally, we assigned spatially resolved transcripts to individual cell bins using boundaries defined by Voronoi tessellation seeded at the transformed nuclear coordinates (doi.org/10.1101/2024.11.07.622434). Cell bins that passed quality-control criteria based on transcript count, detected gene count, and segmentation quality were retained for downstream analyses. Through this procedure, we assigned spatially resolved transcripts to segmented cells and generated cell-bin-level gene expression profiles.

#### Registration to spinal cord atlas

We assigned each cell bin to one of 71 annotated anatomical regions based on histological landmarks and experimental metadata. Anatomical registration was performed in three steps. First, we determined the spinal segment of each sequenced section by examining the adjacent Nissl-stained section. We compared multiple anatomical landmarks, such as overall spinal cord morphology, the boundary between white and grey matter, the size and location of motor neuron groups, sympathetic and parasympathetic neurons, together with the serial sectioning order, against the P56 Allen Mouse Spinal Cord Atlas (https://mousespinal.brain-map.org; Watson et al.). Using these features, we assigned each spinal cord hemisphere to the corresponding atlas segment. We further cross-validated segment assignments using the RNA expression patterns of a curated panel of genes diagnostic of rostrocaudal spinal levels.

Second, we aligned a vectorized version of the Allen Mouse Spinal Cord Atlas to UMI spatial maps reconstructed from the Stereo-seq data. We customized the vectorized atlas from the original reference by assigning a unique colour code to each anatomical region, which enabled subsequent mapping of cell bins to individual regions. Notably, these UMI maps revealed detailed cytoarchitectural features resembling those observed in Nissl-stained sections. This correspondence likely reflects regional differences in cell density between grey and white matter, as well as spatial variation in overall RNA abundance captured by Stereo-seq. We then registered the vectorized atlas to the corresponding UMI spatial map in Adobe Illustrator by globally warping the atlas while preserving the relative positions, shapes, and topology of the annotated spinal cord regions.

Finally, we assigned anatomical identities to segmented cell bins using custom MATLAB scripts. Because the registered atlas and segmented nuclei shared the same coordinate system, each cellular nucleus could be mapped to an anatomical region based on its spatial coordinates and the corresponding region-specific pixel colour in the registered atlas. Through this procedure, each segmented cell bin was assigned to a unique anatomical region.

The regions include: 1Sp: lamina 1 of the spinal grey; 2SpI: lamina 2 of the spinal grey, inner part; 2SpO: lamina 2 of the spinal grey, outer part; 3Sp: lamina 3 of the spinal grey; 4Sp: lamina 4 of the spinal grey; 5Sp: lamina 5 of the spinal grey; 5SpL: lamina 5 of the spinal grey, lateral part; 5SpM: lamina 5 of the spinal grey, medial part; 6Sp: lamina 6 of the spinal grey; 6SpL: lamina 6 of the spinal grey, lateral part; 6SpM: lamina 6 of the spinal grey, medial part; 7Sp: lamina 7 of the spinal grey; 8Sp: lamina 8 of the spinal grey; 10Sp: lamina 10 of the spinal grey; Ad9: adductor motoneurons of lamina 9; asp: anterior spinal artery; Ax9: axial muscle motoneurons of lamina 9; Bi9: biceps motoneurons of lamina 9; cc: central canal; CeCv: central cervical nucleus; CEx9: crural extensor motoneurons of lamina 9; CFl9: crural flexor motoneurons of lamina 9; Cr9: cremaster motoneurons of lamina 9; cu: cuneate fasciculus; D: dorsal nucleus (Clarke); dcs: dorsal corticospinal tract; De9: deltoid motoneurons of lamina 9; dr: dorsal root; ExA9: external anal sphincter motoneurons of lamina 9; ExU9: external urethral sphincter motoneurons of lamina 9; FA9: external urethral sphincter motoneurons of lamina 9; FEx9: forearm extensor motoneurons of lamina 9; FFl9: forearm flexor motoneurons of lamina 9; Gl9: gluteal motoneurons of lamina 9; gr: gracile fasciculus; Hm9: hamstring motoneurons of lamina 9; IB: internal basilar nucleus; ICl: intercalated nucleus; ICo9: intercostal muscle motoneurons of lamina 9; IH9: infrahyoid muscle motoneurons of lamina 9; IML: intermediolateral column; IMM: intermediomedial column; LatC: lateral cervical nucleus; LD9: latissimus dorsi motoneurons of lamina 9; LDCom: lumbar dorsal commissural nucleus; lf/vf: lateral funiculus/ventral funiculus; LPrCb: lumbar precerebellar nucleus; LS9: levator scapulae motoneurons of lamina 9; LSp: lateral spinal nucleus; Man9: manus motoneurons of lamina 9; Pec9: pectoral muscle motoneurons of lamina 9; Pes9: pes motoneurons of lamina 9; Ph9: phrenic motoneurons of lamina 9; Ps9: psoas motoneurons of lamina 9; psdc: postsynaptic dorsal column pathway; Q9: quadriceps motoneurons of lamina 9; QL9: quadratus lumborum motoneurons of lamina 9; Rh9: rhomboid muscle motoneurons of lamina 9; rs: rubrospinal tract; SDCom: sacral dorsal commissural nucleus; SI9: supraspinatus and infraspinatus motoneurons of lamina 9; SM9: sternomastoid motoneurons of lamina 9; SPrCb: sacral precerebellar nucleus; SPSy: sacral parasympathetic nucleus; Sr9: serratus anterior motoneurons in lamina 9; Tail9: tail muscle motoneurons of lamina 9; ThAb9: thoracoabdominal wall muscle motoneurons of lamina 9; Tr9: triceps motoneurons of lamina 9; Tz9: trapezius motoneurons of lamina 9; TzSM9: trapezius and sternomastoid motoneurons of lamina 9; vr: ventral root.

For downstream analyses, the muscle-specific motor nuclei within lamina 9 were grouped into a single lamina 9 label. Because these nuclei are anatomically sparse and finely subdivided, individual nuclei contained too few cells for meaningful statistical analysis. Following this consolidation, a total of 37 anatomical labels were used for downstream analyses.

#### Batch correction and integration

Cell-bin matrices from all sections were integrated with Seurat v5 [58]. Because morphology varies across sections and segments, cell-bin coordinates were centred, scaled and reoriented, and all sections (C, T, L, S) were spatially aligned using the central canal as the primary anatomical anchor. To control technical variation across sections, individual count matrices were merged into a single Seurat object with per-segment layers, normalized and variance-stabilized independently with SCTransform [59], and reduced by principal component analysis (PCA) on Pearson residuals. Batch effects were corrected across segments by applying Harmony [60] to the PCA embedding (IntegrateLayers).

### Data analysis

#### Clustering

The first 50 harmonized principal components were used for UMAP visualization [61] and to construct a shared-nearest-neighbour graph (FindNeighbors); Louvain modularity optimization [62] (FindClusters) was run across resolutions 0.1–1.0, and the final resolution was selected on the basis of cluster stability and canonical marker-gene expression, yielding 12 major cell classes and 94 subclusters. Unsupervised clustering used the top 2,000 highly variable genes (HVGs) selected from expression variance alone; spatially variable genes (SVGs) were used only for the region-level spatial-specificity and co-expression-module analyses described below, so that cell-type and subtype identities were defined independently of anatomical position.

#### Cell-type annotation and anatomical assignment

Major classes were annotated using canonical markers and differential expression: excitatory neurons, inhibitory neurons, motor neurons, astrocytes, oligodendrocytes, OPCs, microglia, Schwann cells, meningeal cells, vascular endothelial cells, erythroid cells and central-canal– associated (ependymal) cells. Representative Stereo-seq marker-gene patterns were compared qualitatively with Allen Mouse Spinal Cord Atlas in situ hybridization images for spatial validation of marker localization and major cell-type annotation.

#### Subclustering

Cells of each major class were subset and independently re-normalized, scaled, reduced and clustered by graph-based clustering to resolve subtype-level structure (15 excitatory, 17 inhibitory, 7 motor-neuron and 55 non-neuronal subclusters). Differential expression against the remaining cells of the same class identified representative markers, which were visualized as violin plots and heatmaps. On the basis of shared anatomical distribution, excitatory subclusters were grouped into four spatial cluster sets (G1 superficial, G2 upper-mid, G3 pan-laminar, G4 deep), and inhibitory subclusters into superficial-dorsal, upper-mid-dorsal, pan-laminar and deep groups. Group-and subtype-specific enrichment of neurotransmitter-receptor, transcription-factor and GPCR gene families was assembled into heatmaps [63]. Segment-resolved composition (average cell count and relative proportion per subcluster) was quantified across levels and individual segments.

#### Spatial-enrichment analysis

Anatomical regions within each transverse section were defined using cytoarchitectural landmarks and region-restricted marker genes (superficial dorsal horn, deep dorsal horn, intermediate zone, ventral horn, central canal, white-matter compartments and peripheral territories). For each subcluster, a spatial-enrichment Z-score was computed across regions as the normalized deviation of observed cell density from a spatially uniform expectation. Scores were assembled into a subcluster-by-region matrix, visualized as a heatmap, and ordered by spatial specificity assessed from the variability of enrichment across regions; spatial-specificity groups were defined using spatial-autocorrelation-based metrics.

#### Spatial density and positional analyses

Annotated identities were projected back onto native cell coordinates for each section, and representative C/T/L/S sections were selected to illustrate spatial organization. For cross-section comparison, tissue outlines were registered to a common template by affine transformation on boundary landmarks, and each section was rotated, aligned and centred. Two-dimensional kernel density estimates of cell positions were computed within the standardized coordinate frame to generate class density maps, and one-dimensional kernel density estimates along the mediolateral and dorsoventral axes were used to build ridge plots (median positions indicated).

#### Gene-level spatial-specificity and module analyses

Mean log-normalized expression of 21,293 detected genes was summarized across registered anatomical regions from all 81 samples. Spatially variable genes (SVGs) were identified using a spatial-specificity loss-entropy metric, in which genes with expression concentrated in fewer regions received higher scores; genes were classified as high (5.7%; 1,212 genes), medium (13.8%; 2,941 genes) or low (80.5%; 17,138 genes) specificity (category counts sum to 21,291; the two-gene difference from the 21,293 detected reflects genes without an assigned category). Spatial gene modules were inferred from local gene–gene co-expression using the spatial-module model implemented in the StereoPy framework [64]: local correlations were computed from spatial expression profiles, genes with similar local patterns were grouped into modules, module activity was summarized as spatial Z-scores and projected onto representative transverse sections, and module–region relationships were evaluated against anatomical regions.

#### Cluster-marker genes (CMGs)

To identify cluster-marker genes (CMGs), differential expression analysis was performed across anatomically defined clusters using the FindAllMarkers function in Seurat (minimum expression threshold: 5% of cells per cluster), yielding the top 2,000 CMGs. To assess the spatial consistency of these CMGs across sexes (male versus female) and anatomical segments (C, cervical; T, thoracic; L, lumbar; and S, sacral), CMGs were independently identified within each respective subset. The global CMG set was then intersected with these subset-specific CMG lists, and the degree of conservation was visualized as an UpSet plot across spinal levels (Figure S2E) and a stacked bar chart across sexes (Figure S2F). This marker-based gene set (top 2,000 cluster markers) is distinct from the spatial-specificity (loss-entropy) SVG classification used for the co-expression-module analysis. The comparison between sexes used L4 (5 males and 4 females) and S1 (3 males and 5 females) segments.

#### Cell-type spatial atlas

To build a spatial atlas independent of direct gene-expression clustering, each cell was represented by a neighbourhood-composition vector [65,66] encoding the local abundance or fraction of neighbouring annotated cell types. These vectors were used for dimensionality reduction, UMAP visualization and clustering. Level 1 clusters defined broad spatial-atlas modules, and Level 2 clusters provided finer anatomical subdivisions. Modules were compared with canonical anatomical zones by computing, for each module, the percentage of its cells assigned to each registered anatomical region; the distribution of transcriptomic subclusters across Level 2 modules was computed analogously.

#### Three-dimensional reconstruction

For 3D visualization, aligned 2D section coordinates were combined with rostrocaudal segment order to assign each section a z-position, and cells or spatial-atlas modules were rendered across C1–S4 to reconstruct longitudinal organization along the whole cord.

### Statistics

For reproducibility analysis, SCTransformed gene-expression profiles were aggregated per sample or segment and compared by Pearson correlation for same-segment/cross-animal, cross-segment/same-animal and cross-segment/cross-animal comparisons; summary plots show mean coefficients with standard deviation. The coefficient of determination (R^2^) was also reported for paired expression profiles. Differential-expression, spatial-enrichment and module analysis used the tests, significance thresholds and multiple-testing corrections specified in the analysis scripts. Statistical analyses were performed in R version 4.5.1.

## Data and code availability

The Stereo-seq data have been deposited in Gene Expression Omnibus (GSE348658). Analysis code used to reproduce the results is available on GitHub (https://github.com/jingli-IBT/mouse-spinal-cord-spatial-transcriptomics/).

## Acknowledgments

This work was supported by grants from the U.S. National Institutes of Health (DC008983, DC020887, MH116990 and NS132912 to L.I.Z. and EY019049 and AG089756 to H.W.T.). K.K.Z. was supported by the Texas A&M AgriLife Institute for Advancing Health Through Agriculture (IHA) and the U.S. Department of Agriculture, Agricultural Research Service, under Agreement No. 58-3091-1-018.

## Author contributions

L.I.Z. and K.K.Z. supervised the study. G.W.Z. and J.J.H. generated the Stereo-seq data. G.W.Z., J.L. and J.J.H. processed and analyzed the data. L.I.Z., K.K.Z., H.W.T. and G.W.Z. wrote the manuscript with input from all authors.

---

**Figure S1.**
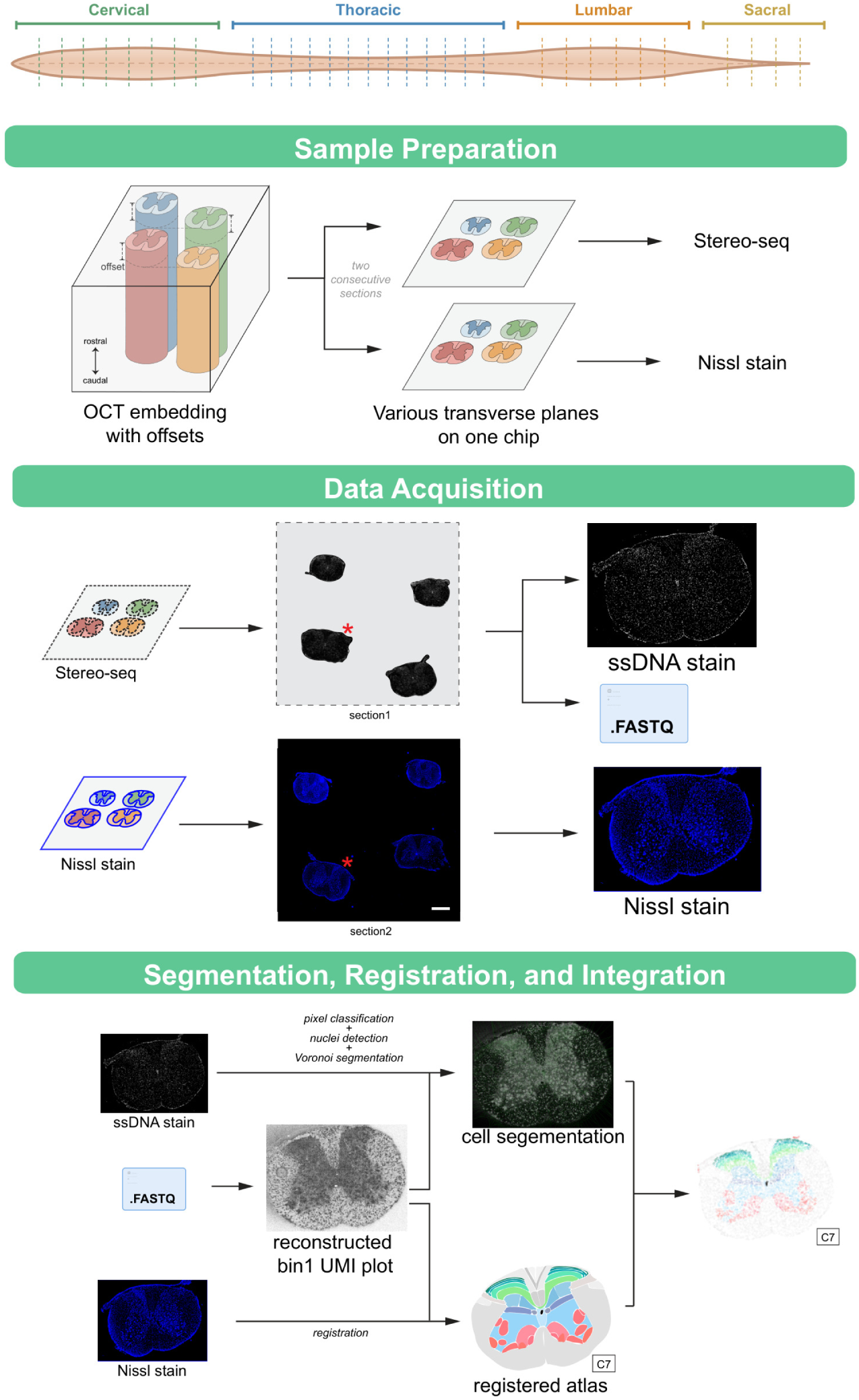
Experimental design and preprocessing workflow. Workflow for whole-spinal-cord spatial transcriptomic profiling from cervical to sacral levels. Multiple spinal cords were co-embedded in OCT with rostrocaudal offsets so that spinal cords captured on a single chip were from different transverse planes. Consecutive sections processed for Stereo-seq and Nissl staining. Stereo-seq procedure generated single-stranded DNA (ssDNA)-stained images and spatially resolved transcriptomic data, which were processed through pixel classification, nuclei detection, Voronoi segmentation and reconstruction of transcript-containing cellbins. Nissl staining provided histological information for registration of reconstructed spatial maps to a spinal cord reference atlas, integrating transcriptomic, spatial and anatomical information for each segmented cellbin.

**Figure S2.**
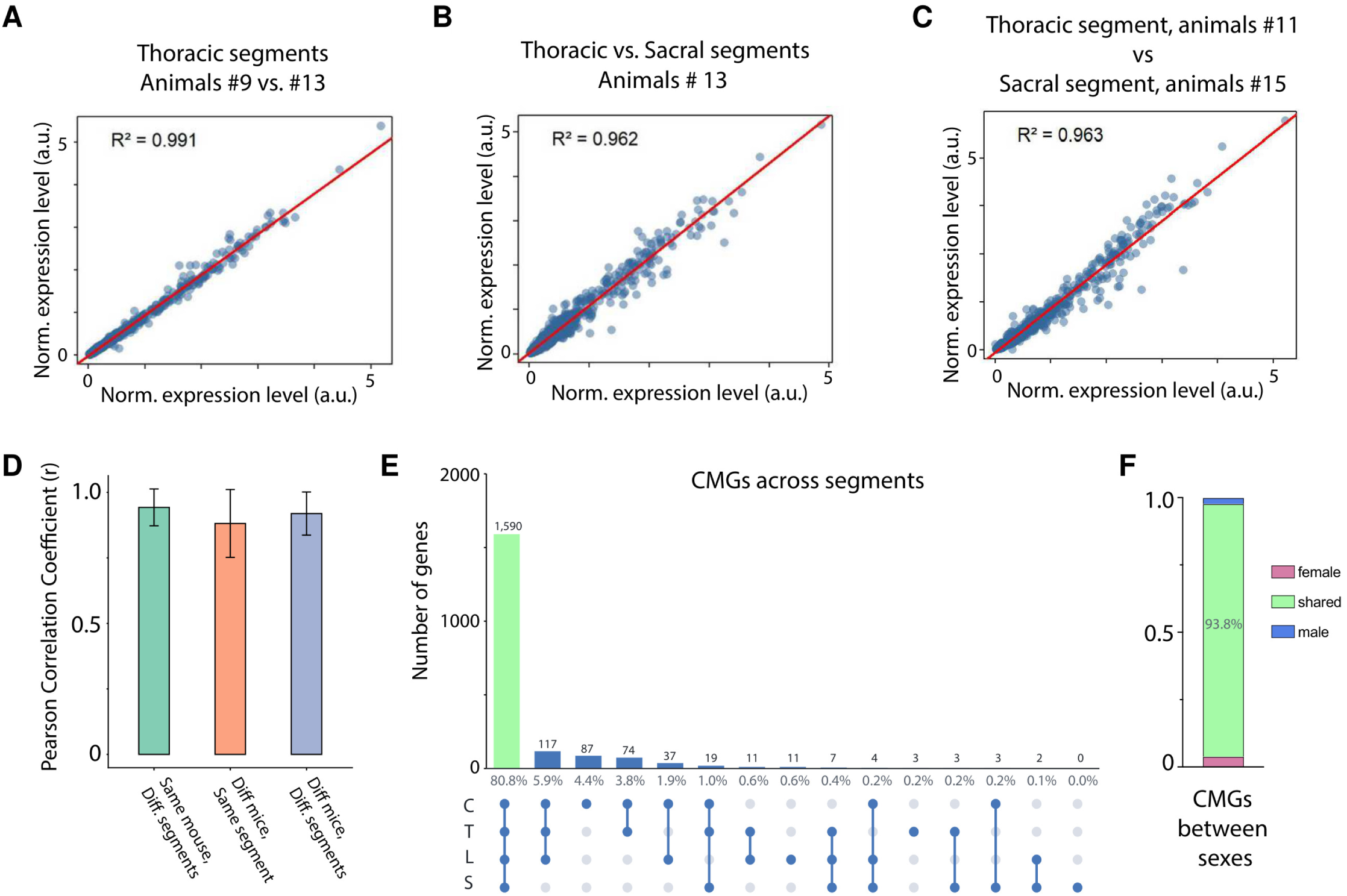
Quality control and reproducibility assessment of spatial transcriptomic data. A-C. Scatter plots showing correlations of SCTransformed gene expression values across mice and anatomical segments. A. Biological reproducibility: Correlation between thoracic segments from animals #9 and #13 (R^2^ = 0.991), showing high similarity between different mice within the same spinal segment. B. Inter-segment comparison: Correlation between thoracic and sacral segments from animal #13 (R^2^ = 0.962), showing transcriptomic similarity between different anatomical segments within the same mouse. C. Cross-animal segment comparison: Correlation between a thoracic segment from animal #11 and a sacral segment from animal #15 (R^2^ = 0.963), showing high similarity across different animals and different spinal segments. Note: All axes represent SCTransformed expression values. The red line indicates the identity line (y = x). D. Bar plots of the Pearson correlation coefficient (r) summarizing pairwise comparisons across Same mouse/Diff. segment, Diff. mouse/Same segment and Diff. mouse/Diff. segment categories. Data are mean + s.d. E. UpSet plot showing the number of spatially variable genes(SVGs) unique to, or shared among, cervical, thoracic, lumbar and sacral sections. Counts are shown above each bar and percentages below; the set shared across all four levels is highlighted in green. F. Stacked bar chart showing spatially variable genes(SVGs) shared between, or unique to, male and female sections (1,940 SVGs total: 93.8% shared, 3.9% female-specific, 2.3% male-specific).

**Figure S3.**
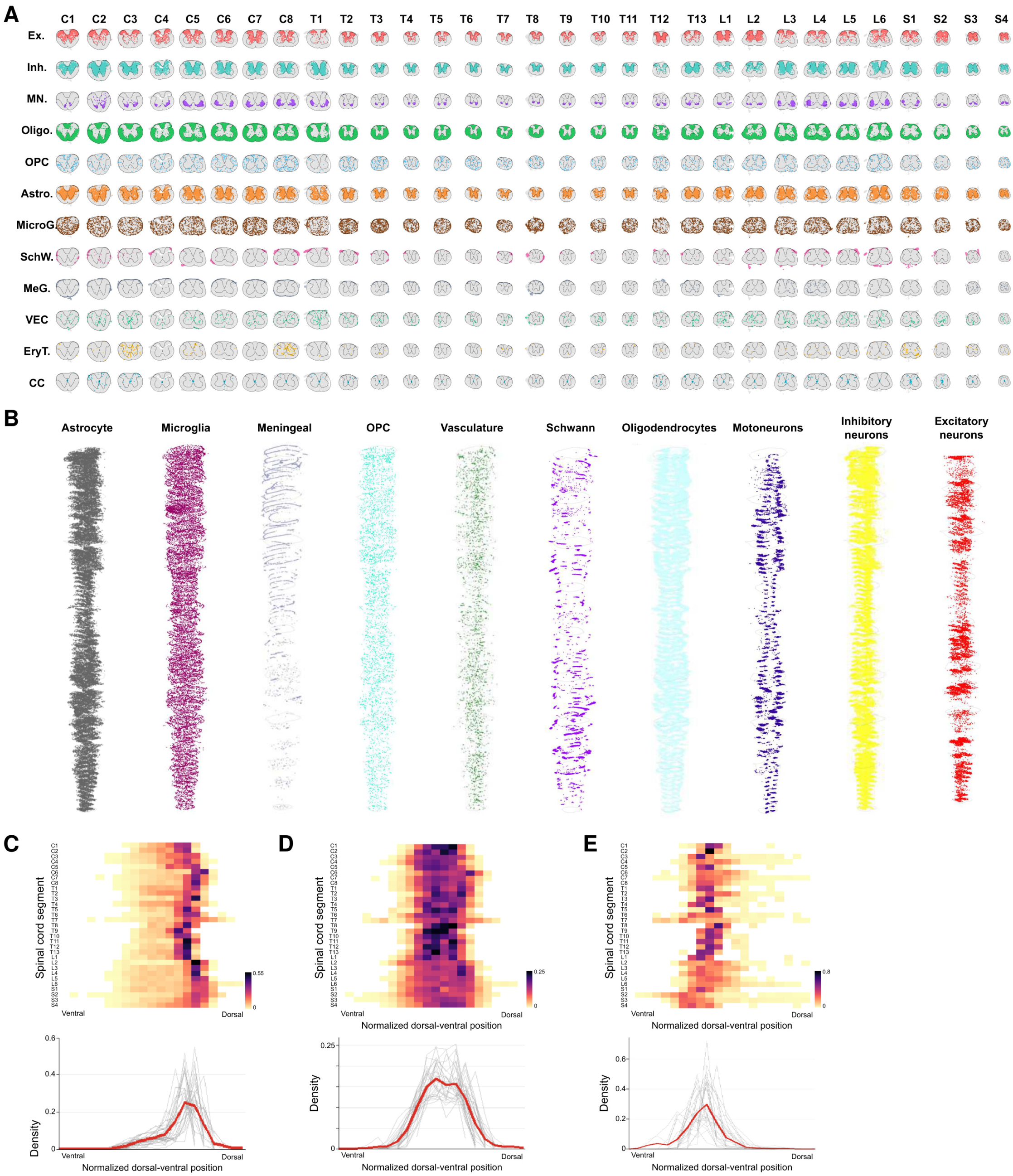
Spatial distribution and density profiles of neuronal and non-neuronal cell types. A. 2D spatial mapping of major clusters: Spatial distributions of major cell classes across all spinal segments (C1-C8, T1-T13, L1-L6, S1-S4), shown as one row per major cell class. Each class shows a distinct anatomical preference within the white or grey matter. B. 3D spinal cord reconstruction: Longitudinal 3D visualizations of individual major cell classes along the spinal cord axis, shown in separate columns (astrocytes, microglia, meningeal cells, OPCs, vasculature, glia, Schwann cells, oligodendrocytes, motoneurons, inhibitory neurons and excitatory neurons). C-E. Normalized spatial density and consistency profiles: Quantitative analysis of cell distribution relative to the dorsal-ventral axis. Heatmaps (top): Cell density for selected major cell classes across spinal segments, normalized along the dorsal-ventral axis. The colour scale indicates cell density and highlights segment-wise spatial positioning. Density profiles (bottom): Overlaid spatial consistency plots across spinal segments. Grey lines represent individual segment profiles, and the red line indicates the mean density profile, showing conservation of spatial positioning along the spinal cord.

**Figure S4.**
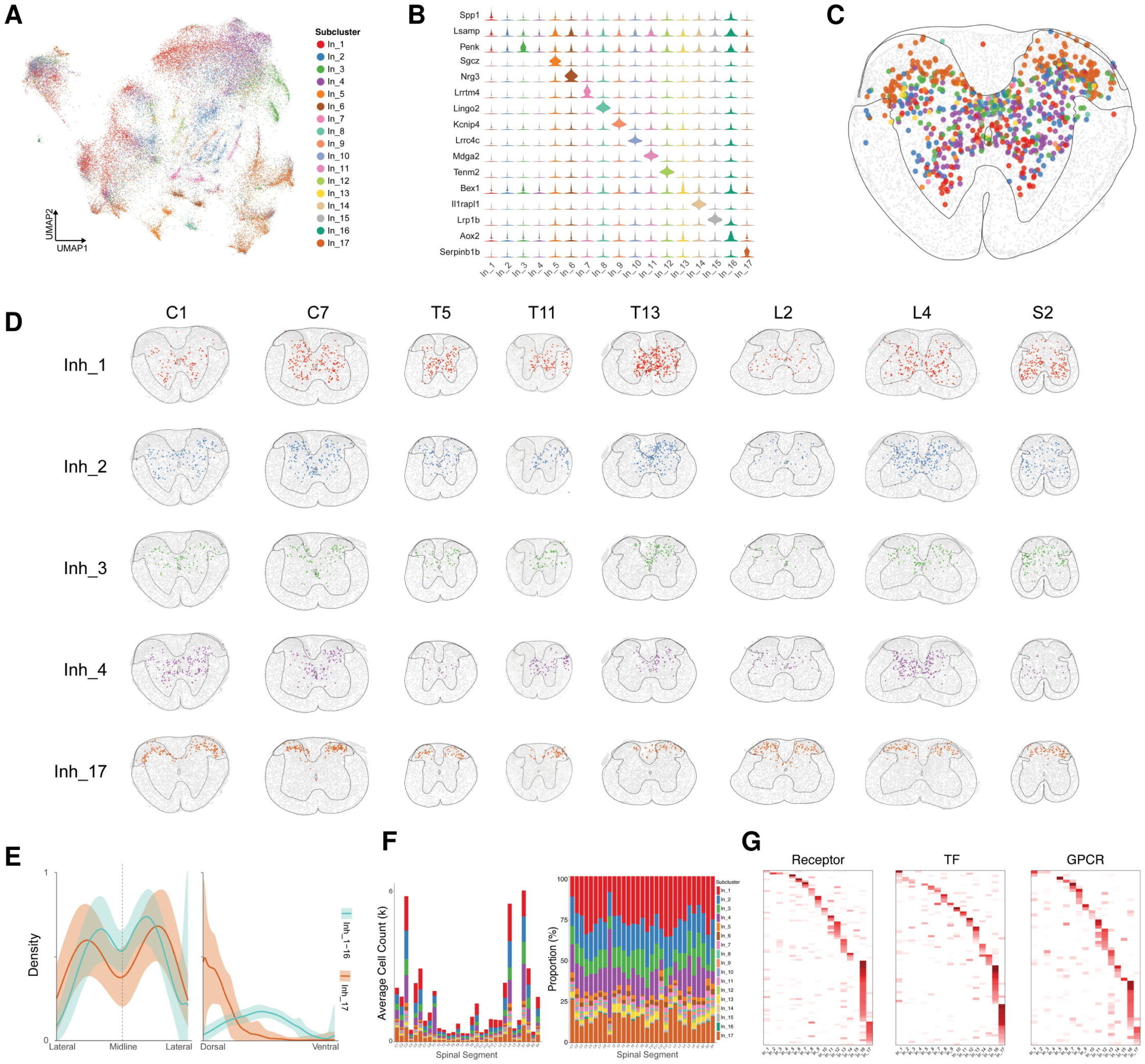
Spatial organization of inhibitory neuron subtypes across the whole mouse spinal cord. A. UMAP of inhibitory neurons coloured by subcluster (Inh_1-Inh_17). B. Violin plots of representative markers across subclusters. C. Representative transverse section showing the spatial distribution of inhibitory subclusters. D. Spatial distributions of selected groups (Inh_1, Inh_2, Inh_3, Inh_4, Inh_17) across representative spinal segments. E. Density plots comparing mediolateral and dorsoventral distributions of Inh_17 vs other inhibitory subclusters. F. Average cell count (left) and proportional composition (right) of subclusters across segments. G. Heatmaps of receptor, transcription-factor and GPCR gene expression across inhibitory subclusters.

**Figure S5.**
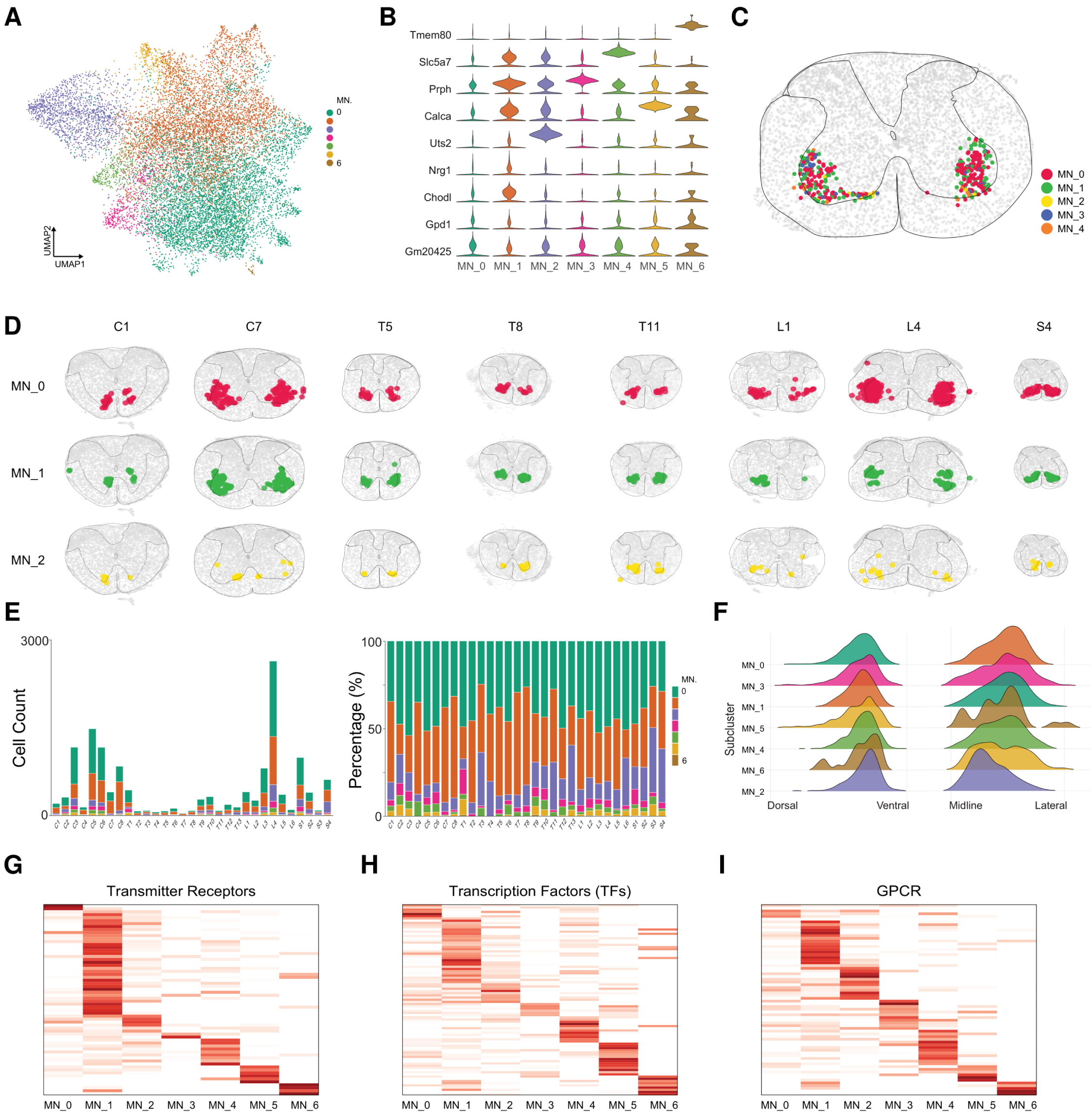
Spatial organization of motor neuron subtypes across the whole mouse spinal cord. A. UMAP of motor neurons coloured by subcluster (MN_0-MN_6). B. Violin plots of representative markers, including Slc5a7, Prph and Chodl and subtype-specific genes. C. Representative transverse section showing selected motor-neuron subclusters (MN_0-MN_2) in the ventral horn. D. Spatial distributions of MN_0-MN_2 across representative spinal segments. E. Cell counts and proportional composition of subclusters across segments. F. Ridge plots of subclusters along the dorsoventral and mediolateral axes. G-I. Heatmaps of transmitter-receptor (G), transcription-factor (H) and GPCR (I) gene expression across motor-neuron subclusters.

**Figure S6.**
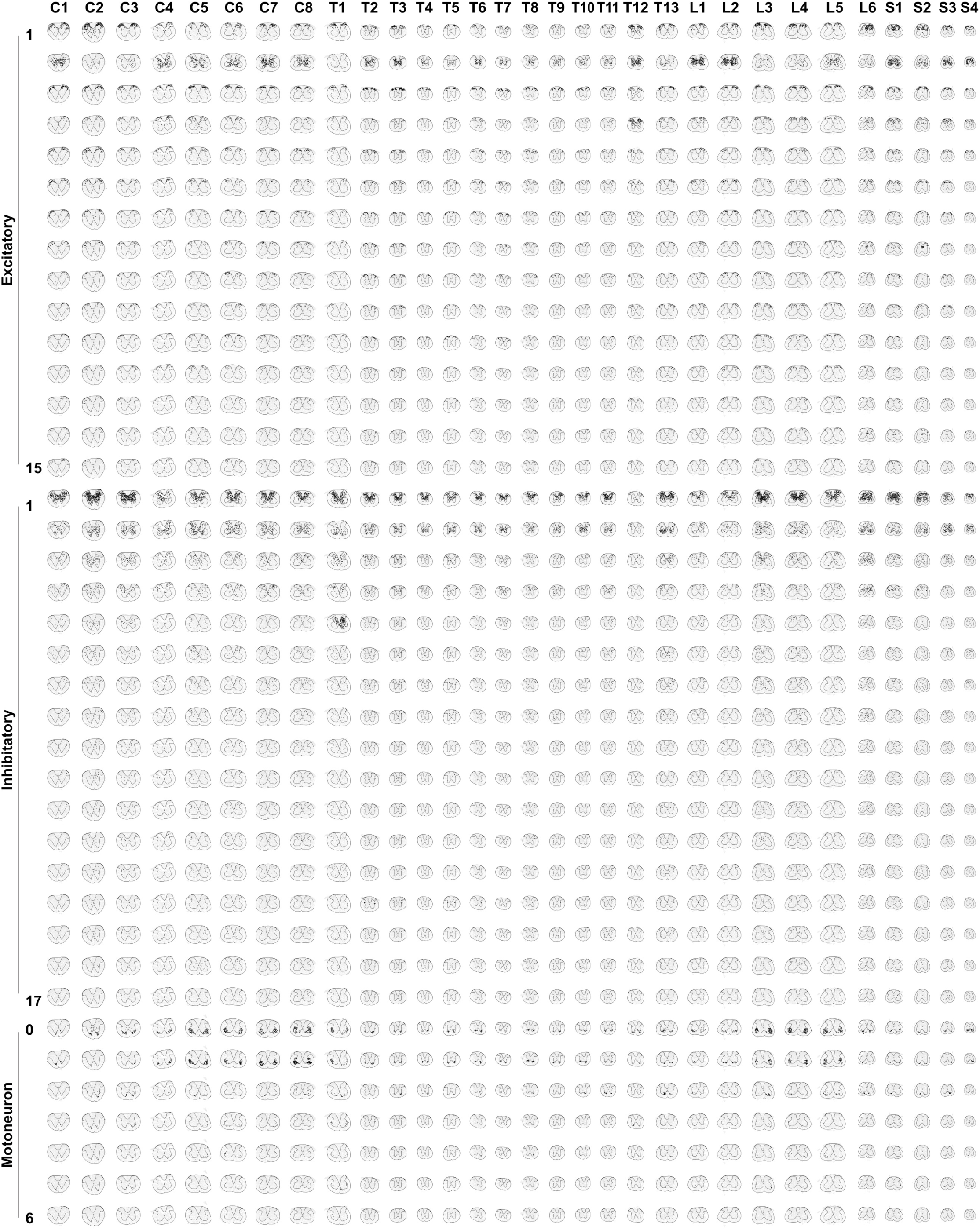
Spatial distribution of neuronal subclusters. 2D Spatial Mapping of Sub-clusters: Spatial distributions of the 94 transcriptomic subclusters, grouped by their 12 major cell classes (excitatory neurons, inhibitory neurons, motoneurons, oligodendrocytes, OPCs, astrocytes, microglia, Schwann cells, meningeal cells, vascular endothelial cells, erythroid cells and central-canal-associated cells). Rows show individual subclusters within each neuronal major class; columns are spinal segments from C1 to S4, with alternate segments labelled. Each subcluster is projected onto transverse spinal cord sections, revealing distinct and reproducible spatial territories across the rostrocaudal axis. Glial and other non-neuronal classes are shown in Figures S7 and S8.

**Figure S7.**
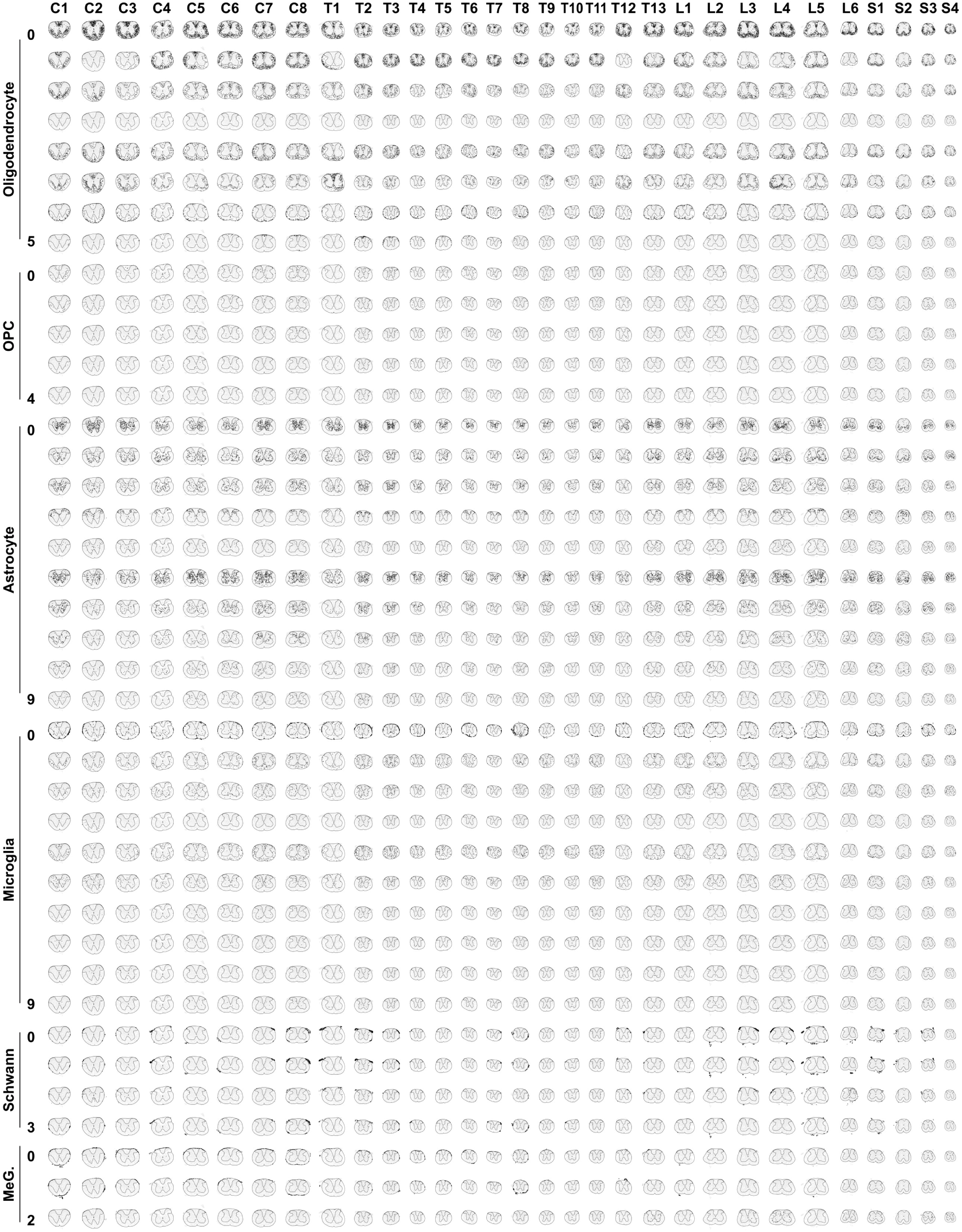
Spatial distribution of glial subclusters. Rows show individual glial subclusters within each major class; columns are spinal segments from C1 to S4, with alternate segments labelled. Each subcluster is projected onto transverse spinal cord sections, revealing distinct and reproducible spatial territories across the rostrocaudal axis.

**Figure S8.**
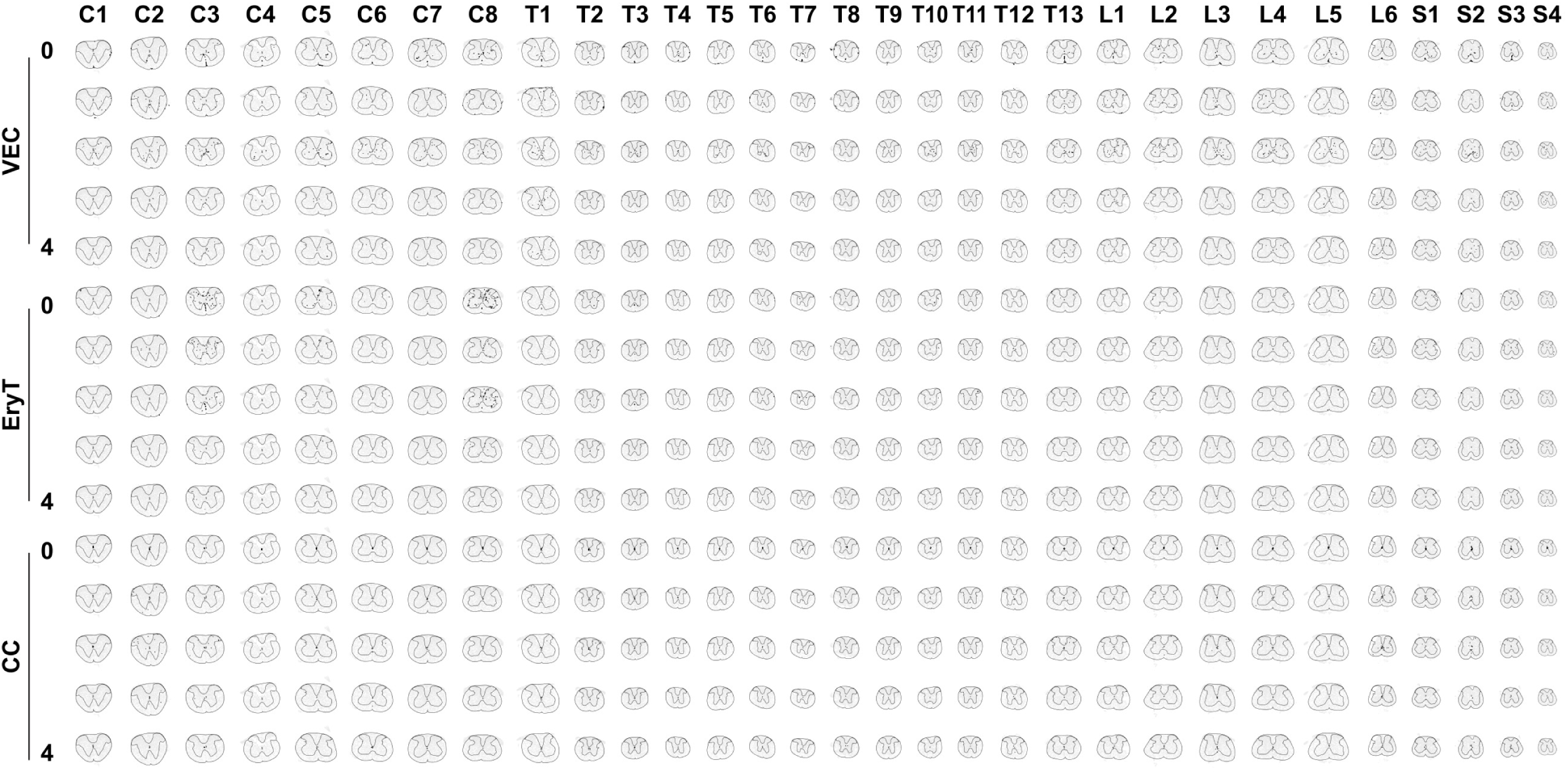
Spatial distribution of other non-neuronal subclusters. Rows show individual non-neuronal subclusters within each major class; columns are spinal segments from Cl to S4, with alternate segments labelled. Each subcluster is projected onto transverse spinal cord sections, revealing distinct and reproducible spatial territories across the rostrocaudal axis.

